# Zebrafish Polycomb repressive complex-2 critical roles are largely Ezh2- over Ezh1-driven and concentrate during early embryogenesis

**DOI:** 10.1101/2020.12.31.424918

**Authors:** Gabriel A. Yette, Scott Stewart, Kryn Stankunas

## Abstract

Polycomb repressive complex-2 (PRC2) methylation of histone H3 lysine-27 (H3K27me) is associated with stable transcriptional repression. PRC2 famously silences *Hox* genes to maintain anterior-posterior segment identities but also enables early cell fate specification, restrains progenitor cell differentiation, and canalizes cell identities. Zebrafish PRC2 genetic studies have focused on *ezh2,* which, with its paralog *ezh1*, encodes the H3K27 methyltransferase component. *ezh2* loss-of-function mutants reinforce essential vertebrate PRC2 functions during early embryogenesis albeit with limited contributions to body plan establishment. However, redundancy with *ezh1* and the lethality of maternal-zygotic homozygous *ezh2* nulls could obscure additional early developmental and organogenesis roles of PRC2. Here, we combine new and existing zebrafish *ezh1* and *ezh2* alleles to show collective maternal/zygotic *ezh2* exclusively provides earliest embryonic PRC2 H3K27me3 activity. Zygotic *ezh1*, which becomes progressively expressed as *ezh2* levels dissipate, has minor redundant and noncompensatory larval roles but itself is not required for viability or fertility. Zygotic Ezh2/PRC2 promotes correct craniofacial bone shape and size by maintaining proliferative pre-osteoblast pools. An *ezh2* allelic series including disrupted maternal *ezh2* uncovers axial skeleton homeotic transformations and pleiotropic organogenesis defects. Further, once past a critical early window, we show zebrafish can develop near normally with minimal bulk H3K27me3. Our results suggest Ezh2-containing PRC2 stabilizes rather than instructs early developmental decisions while broadly contributing to organ size and embellishment.

## INTRODUCTION

Polycomb Repressive Complex 2 (PRC2) directs histone H3 lysine 27 (H3K27me) methylation associated with Polycomb group (PcG)-mediated transcriptional repression (Ferrari et al., 2014). PcG was originally characterized in *Drosophila* for its conserved role maintaining Hox expression patterns and therefore segment identity (Lewis, 1978; reviewed in Kassis et al., 2017). Vertebrate studies, largely using mice, show PRC2 has diverse roles in body plan establishment, cell fate specification, identity maintenance, and growth control during organogenesis and homeostasis (reviewed in Laugesen & Helin, 2014; Scelfo et al., 2015). However, a unifying model how vertebrate PRC2 integrates into developmental regulatory networks remains elusive. Zebrafish provide an additional model organism to explore core vertebrate PRC2 functions. Genetic perturbation of zebrafish *ezh2,* the predominant PRC2 methyltransferase, suggest less essential roles for PRC2 in early body patterning than expected from mouse studies (San et al., 2016). However, a potentially redundant role for the alternative PRC2 methyltransferase *ezh1* is unexplored. Likewise, PRC2 roles in zebrafish organ development and maintenance and how those compare with mammals are largely unknown.

Vertebrate PRC2 comprises four core proteins, with subunits Ezh1 or Ezh2 as H3K27 methyltransferases. PRC2 deposits all mono, di, or tri-methylation H3K27 marks with H3K27me3 at regulatory sites most associated with PRC2-repressed genes (Cao et al., 2002; Müller et al., 2002; Ferrari et al., 2014; Laugesen et al., 2019). PRC2/H3K27me3 cooperates with PRC1 and its histone H2A mono-ubiquitin lysine 119 catalyzed marks to promote stable gene repression (reviewed in Di Croce & Helin, 2013; Wang et al., 2004). H3K27me3 genomic landscapes vary between cells of different identities and states, largely correlating with repression. Therefore, recruited PRC2 activity may promote expression program dynamics underlying cell transitions and/or reinforce transiently induced repression into stable “silencing” that maintains cell fates and states (reviewed in Scelfo et al., 2015; Yu et al., 2019). Conversely, H3K27me3 removal by Kdm6 H3K27me3 demethylases or nucleosome turnover could enable the timely reversal of otherwise stable cell conditions (Chory et al., 2019; reviewed in Gökbuget & Blelloch, 2019; Kartikasari et al., 2013; Li et al., 2014; Rocha-Viegas et al., 2014; Seenundun et al., 2010). Regardless, PRC2-mediated silencing seemingly provides “epigenetic” memory at specific loci to anchor cell decisions and/or more generally buffers against spurious transcriptional noise. Comparing PRC2 roles in diverse developmental contexts facilitates a core understanding how it integrates into regulatory networks including whether PRC2 instructs vs. maintains transcriptional states and if H3K27me3 repression is obligate or supportive.

Mammalian PRC2 studies link its function to early cell fate specification. Many fate-determining genes in embryonic stem (ES) cells exhibit a “bivalent state” of both H3K27me3 and its opposing H3K4me3 mark deposited by Trithorax group/COMPASS complexes (Bernstein et al., 2006). Upon lineage commitment, the antagonistic marks resolve in favor of either stable repression (H3K27me3) or activation (H3K4me3). Accordingly, ES cells largely fail to differentiate in the absence of PRC2 (Chamberlain et al., 2008; Ferrari et al., 2014; Lavarone et al., 2019; Pasini et al., 2007). *Ezh2-null* mice, and mutants in other PRC2 components, show early embryonic lethality around gastrulation, consistent with PRC2 contributing to the earliest cell specification decisions (Faust et al., 1995; O’Carroll et al., 2001; Pasini et al., 2004).

Vertebrate PRC2 remains essential for organogenesis after the initial body plan and organ specific cell lineages are defined. In particular, Cre/loxP-mediated cell type specific loss of mouse PRC2 link its functions to maintaining tissue-specific progenitor pools and/or robust cell identities in disparate organs (Chiacchiera et al., 2016; Delgado-Olguín et al., 2012; Dudakovic et al., 2015; Ezhkova et al., 2011; Koppens et al., 2016; Snitow et al., 2015). However, varying conclusions from in vivo and cultured cell experiments could be clarified and expanded by comparative evaluation of PRC2 functions in another vertebrate model. For example, in skeletal development, a focus herein, mouse *Ezh2* studies variably link PRC2/H3K27me3 to specification and expansion of osteoblast precursor population, osteoblast progenitor maintenance and proliferation, and differentiation into mature, bone-producing osteoblast state (Dudakovic et al., 2015; Ferguson et al, 2018; Lui et al., 2016; Mirzamohammadi et al., 2016; Schwarz et al., 2014).

Zebrafish provide a compelling second vertebrate model to explore PRC2 roles. H3K27me3 marks are first detected at zygotic genome activation (ZGA; 3 hours post fertilization) preceding gastrulation (Lindeman et al., 2010; Murphy et al., 2018; Vastenhouw et al., 2010; Zhang et al., 2014). Homozygous *ezh2* zygotic zebrafish mutants survive to around 12 days post fertilization (hpf) before succumbing to organogenesis defects of the gut and presumably other systems (Dupret et al., 2017; San et al., 2018). Maternal *ezh2* is dispensable for early zebrafish embryogenesis (San et al., 2016). However, maternal zygotic *ezh2* mutants die around two days post fertilization with defects in eye, brain, heart and gut development. Nevertheless, they still establish a normal body plan despite greatly reduced bulk H3K27me3 (San et al., 2016).

*ezh1* homozygous mutant zebrafish are viable and fertile, as are mouse *Ezh1* mutants (Ezhkova et al., 2011; Völkel et al., 2019). However, Ezh1, which shows context-dependent redundancy with Ezh2 in mice (Ai et al., 2017; Bae et al., 2015; Ezhkova et al., 2011; Koppens et al., 2016; Lui et al., 2016; Mirzamohammadi et al., 2016; Shen et al., 2008), may partially fulfill the PRC2 methyltransferase role during early zebrafish development. While Ezh1 is a less potent H3K27me3-catalyzing enzyme (Margueron et al., 2008), ES cell studies suggest Ezh1 and Ezh2 have partially overlapping functions. Loss of Ezh2 alone largely abolishes global H3K27me3 (Lavarone et al., 2019; Shen et al., 2008). However, the remaining Ezh1-PRC2 can deposit H3K27me3 at nucleating sites but without typical spreading (Lavarone et al., 2019). Deletion of both Ezh1 and Ezh2 eliminates H3K27me3 with concomitant derepression of a relatively modest number of genes (Lavarone et al., 2019).

Here, we generate and characterize null alleles of *ezh1* and *ezh2* to explore their combined roles in zebrafish early development and organogenesis, with a focus on the skeletal system. We find that Ezh2 (of either maternal or zygotic origin), and not Ezh1, almost entirely contributes the bulk H3K27me3 and PRC2 function immediately following ZGA and through embryogenesis. These results underscore that vertebrate PRC2 is dispensable for initial formation of organs from all three germ layers and overall body plan establishment. We show PRC2 maintains dermal bone pre-osteoblasts including their proliferative state to form craniofacial elements of the correct size and shape. We clarify contradictory assessments of a hypomorphic *ezh2* allele and use it for an allelic series unveiling homeotic phenotypes and diverse maternal/zygotic-dependent organogenesis defects upon PRC2 disruption. Collectively, our studies reinforce that vertebrate PRC2 has broad developmental roles. However, its major functions supporting body plan elaboration including early organogenesis are transiently concentrated immediately after zygotic genome activation. PRC2 roles during later stages of organ development may include maintaining progenitor populations for organ size establishment or homeostasis.

## RESULTS

### Ezh2 and not Ezh1 fulfills most PRC2 roles during early embryogenesis

We explored if and how zebrafish *ezh1* and *ezh2* function divergently or redundantly, including by compensatory mechanisms. We first examined *ezh1* and *ezh2* expression from one hour post fertilization (hpf) through five days post fertilization (dpf) by RT-qPCR (Fig. 1A) and whole mount RNA *in situ* hybridization (Fig. 1B-G). *ezh2* was highly expressed during early stages of development compared to *ezh1*, which was undetectable until 24 hpf (Fig. 1A-F). *ezh2* transcripts progressively decreased while those of *ezh1* increased from 24 hpf to 120 hpf (Fig. 1A, D, G). Expression Atlas zebrafish RNA-Seq developmental time course data (White et al., 2017) showed a similar transition from *ezh2* to *ezh1* relative predominance (Fig. S1A). We did not detect maternal *ezh1* mRNA, consistent with most other studies (Chrispijn et al., 2018; San et al., 2018; San et al., 2016; Sun et al., 2008; White et al., 2017), except one report (Völkel et al., 2019). Regardless, overlapping *ezh1* and *ezh2* expression during zebrafish development suggests Ezh1 and Ezh2 could contribute collectively to embryonic/larval PRC2. If so, loss of both PRC2 methyltransferases should enhance deleterious effects of either mutant alone.

**Figure 1.**
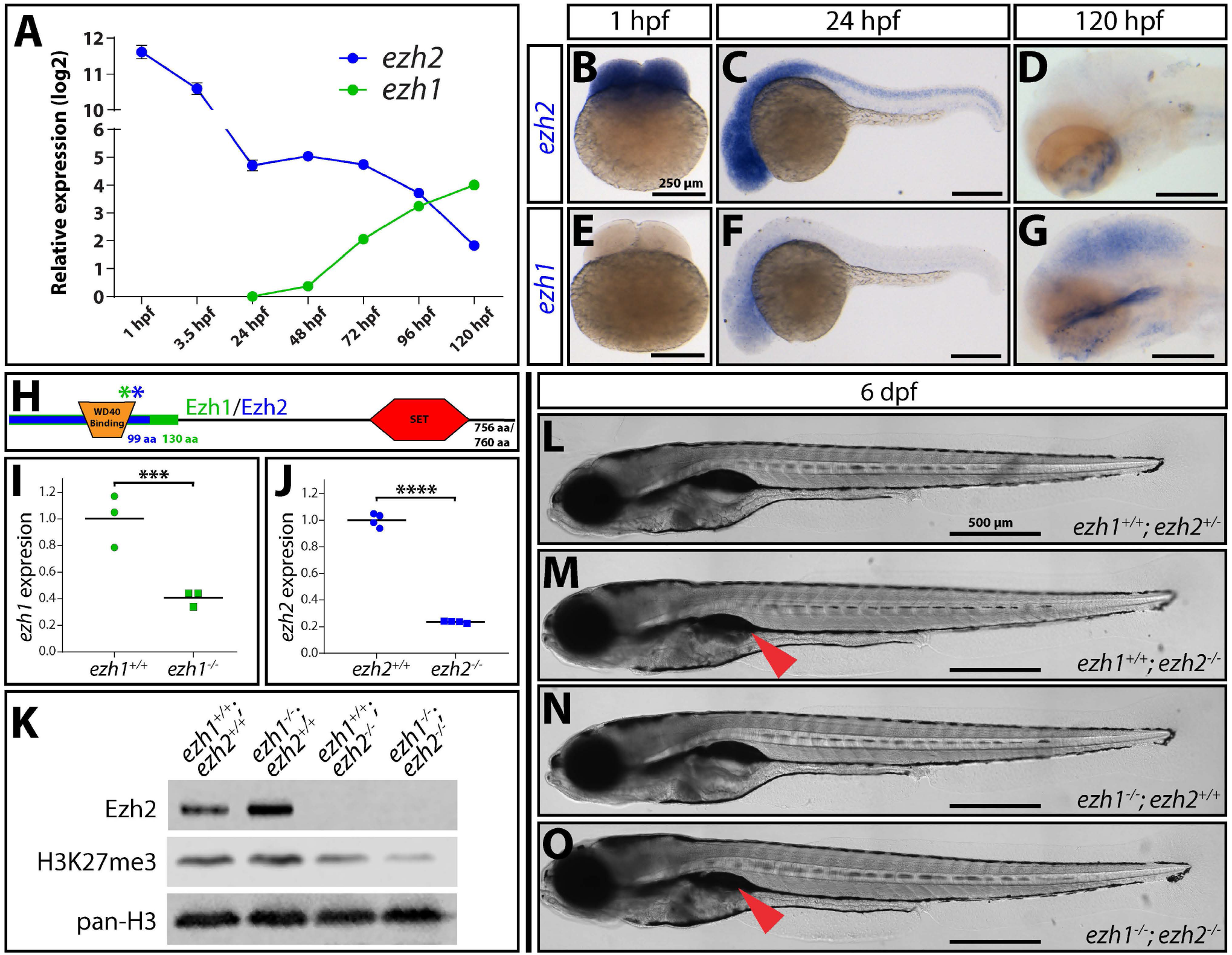
Ezh2 and not Ezh1 fulfills zygotically-produced PRC2’s relatively limited role during zebrafish embryogenesis. **(A)** Graph of RT-qPCR-determined *ezh2* (blue) and *ezh1* (green) relative transcript levels in wildtype embryos/larvae from one to 120 hours post fertilization (hpf). Transcript levels are normalized to *rpl8* (reference gene) and then *ezh1* levels at 24 hpf. **(B-G)** In situ hybridization for *ezh2* (B-D) and *ezh1* (E-G) at 1 hpf, 24 hpf, and 120 hpf. Scale bar = 250 μm. **(H)** Schematic of Ezh1/Ezh2 proteins (756 and 760 amino acids (aa), respectively) with asterisks denoting location of CRISPR/Cas9-generated mutations in new loss-of-function *ezh1* and *ezh2* alleles. Overlaid boxes show predicted truncated polypeptides from the *ezh1* (green, 130 aa), and *ezh2* (blue, 99 aa) alleles. Both mutations disrupt the reading frame and lead to early termination. **(I-J)** Scatter plots comparing (I) *ezh1* transcripts between wildtype and *ezh1* homozygous mutant clutchmates at three dpf and (J) *ezh2* transcripts between *ezh2* mutant and wildtype two day old siblings. RT-qPCR levels are normalized to the means of wildtype groups. *** = P-value < 0.005, **** = P-value < 0.0001 by Student’s two-tailed *t*-tests. **(K)** Immunoblots of whole cell lysates from pooled 6 days post fertilization (dpf) clutchmates using antibodies against Ezh2, H3K27me3, and pan-histone H3. While there is marked decrease in H3K27me3 in *ezh2^−/−^* mutants, notable H3K27me3 remains in *ezh2^−/−^; ezh1^−/−^* samples. **(L-O)** Whole mount differential interference contrast (DIC) microscopy images of 6 dpf control, *ezh1, ezh2*, and *ezh1/ezh2* double homozygous mutant larvae. Red arrowheads indicate under-inflated swim bladder. Scale bar = 500 μm.

We generated *ezh1* and *ezh2* mutants by CRISPR/Cas9-targeting within or downstream of the coding region for the essential N-terminal WD40 binding domain (Fig. 1H). We isolated an *ezh1* allele, *ezh1^b1394^*, missing two bases and an *ezh2* allele, *ezh2^b1393^,* with 7 bases deleted (henceforth referred to as *ezh1^−^* and *ezh2^−^*, respectively) (Fig. S1B, C). Each mutation produced frameshifts predicting early termination after residue 130 and residue 99 of Ezh1 and Ezh2, respectively. The new stop codons precede the catalytic SET domains and therefore likely produce null alleles (Fig. 1H). By RT-qPCR, *ezh1^−/−^* and *ezh2^−/−^* embryos showed 40% and 25% transcript levels for the respective targeted gene than wildtype clutchmates (Fig. 1I-J). The decreased expression likely represented nonsense-mediated decay (NMD) of each mutant transcript, supporting both alleles provide strong loss-of-function. Further, we did not detect any Ezh2 protein in 6 dpf *ezh2^−/−^* larval extracts, reinforcing *ezh2^b1392^* is a null allele (Fig. 1K). Zebrafish mutant alleles can trigger “transcriptional adaptation”, whereby NMD-produced fragments trigger upregulation of related genes that replace lost gene function (El-Brolosy et al., 2019; El-Brolosy & Stainier, 2017; Rossi et al., 2015; Williams et al., 2018). However, neither *ezh1* nor *ezh2* alleles produced compensatory expression of the corollary transcript (Fig. S1D, E). We conclude both alleles likely provide a clean loss-of-function of the respective gene and therefore double homozygous mutants likely eliminate PRC2 function and H3K27 methylation.

We explored relative contributions of Ezh1 and Ezh2 to embryogenesis and H3K27me3 levels by intercrossing *ezh1^+/-^; ezh2^+/-^* fish and examining progeny of all genetic combinations. Loss of Ezh1 had no measurable effect on bulk H3K27me3 in 6 dpf larvae detected by Western blotting (Fig. 1K). Consistently, and matching a previous study (Völkel et al., 2019), *ezh1^−/−^* 6 dpf larvae were indistinguishable from wildtype clutchmates and developed into viable and fertile adults (Fig. 1L, N). Unlike *ezh1* mutants, *ezh2^−/−^* 6 dpf larvae had markedly decreased bulk H3K27me3 (Fig. 1K) and died between 7 and 14 dpf, similar to other *ezh2* null alleles (Dupret et al., 2017; San et al., 2018; San et al., 2016). In addition, *ezh2^−/−^* 6 dpf larvae had underinflated swim bladders (Fig. 1M, O), a phenotype also apparent in published images of homozygotes for other *ezh2* alleles (Dupret et al., 2017; San et al., 2018).

We examined *ezh1; ezh2* double homozygous mutants to define combined Ezh1 and Ezh2 developmental roles. Western blots showed *ezh1^−/−^; ezh2^−/−^* 6 dpf larvae had further reduced bulk H3K27me3 compared to loss of *ezh2* alone (Fig. 1K). This confirmed the loss-of-function nature of the *ezh1* allele while demonstrating bulk H3K27me3 at 6 dpf represented the combined activity of Ezh1- and Ezh2-containing PRC2 complexes. However, *ezh1^−/−^; ezh2^−/−^* larvae still retained some H3K27me3 (Fig. 1K). Given the zebrafish genome includes no other H3K27me3 methyltransferases and maternal or zygotic Ezh2 can fulfill PRC2 function before 2 dpf (San et al., 2016), we infer the residual H3K27me3 likely reflects H3K27me3 deposited by maternal Ezh2 in thereafter slowly dividing or quiescent cells.

We assessed cell type-specific H3K27me3 dependency on Ezh1- and Ezh2-containing PRC2 by immunostaining sectioned wildtype, *ezh1, ezh2,* and *ezh1/ezh2* homozygous mutant 6 dpf larvae. In *ezh1^−/−^* larvae, nuclear H3K27me3 staining was modestly lower in cardiomyocytes and chondrocytes and more markedly reduced, but not absent, in eye, intestine, and central nervous system (CNS) tissue (hindbrain / spinal cord) (Fig. S2, Fig. S3). In contrast, H3K27me3 levels in *ezh1^−/−^* skeletal muscle cells were comparable to wildtype animals (Fig. S3A). H3K27me3 was reduced to a greater extent in *ezh2* zygotic-null animals. H3K27me3 was undetectable in heart, eye, cartilage and gut cells (Fig. S2) and at low levels in skeletal muscle and CNS tissue (Fig. S3). Combined loss of zygotice *ezh1* and *ezh2* further reduced H3K27me3, remaining evident only in select skeletal muscle and CNS cells (Fig. S3). Their minimal H3K27me3 turnover and/or dilution suggests these are quiescent cells, such as satellite cells or radial glia, set aside early in development. Nonetheless, we conclude zygotic Ezh2/PRC2 is responsible for most H3K27me3 in larval zebrafish.

*ezh1^−/−^; ezh2^−/−^* 6 dpf larvae were outwardly indistinguishable from *ezh1^+/+^; ezh2^−/−^* clutchmates despite the former’s even lower H3K27me3 levels (Fig. 1L, M). We crossed *ezh1^−/−^; ezh2^+/-^* females to *ezh1^+/−^; ezh2^+/−^* males to rule out any maternal Ezh1 contribution overlooked by expression studies. Larval and adult (where viable) phenotypes of all given genotypes exactly matched those generated from an *ezh1^+/−^; ezh2^+/−^* in-cross (Fig. S1F-I). These results confirm that Ezh1 does not have a notable maternal role. We additionally conclude Ezh2 provides all phenotypically-evident PRC2/H3K27me3 activity up to 6 dpf. Further, these H3K27me3 deposition roles are concentrated during a short window of development following zygotic genome activation when *either* maternal or zygotic Ezh2 pools can fulfill PRC2’s overt roles (San et al., 2016). Ezh1’s essential roles after its expression increases post-embryogenesis either are masked by low levels of redundant Ezh2 or are outwardly minimal in a laboratory-controlled environment.

### PRC2 promotes craniofacial dermal bone shape and size by maintaining progenitor cell characteristics

Conditional PRC2 mouse studies implicate H3K27me3 dynamics in osteogenesis and skeletal patterning (Dudakovic et al., 2015; Ferguson et al., 2018; Lui et al., 2016; Mirzamohammadi et al., 2016; Schwarz et al., 2014). In zebrafish, we found H3K27me3 demethylases Kdm6ba and Kdm6bb promote expansion of the basihyal craniofacial cartilage (Akerberg et al., 2017). Otherwise, H3K27me3 in the zebrafish skeletal system has not been well characterized. We examined development of the craniofacial skeleton, which comprises endochondral and intramembranous dermal bones (Fig. 2A), in zygotic mutants of *ezh1* and/or *ezh2*. Flat mount skeletal preparations of pharyngeal arch 1 and pharyngeal arch 2 skeletal elements showed that 6 dpf *ezh1^−/−^* larvae had small and malformed osteoblast-derived dermal bones (the opercle and branchiostegal rays (BSR) 2 and 3) (Fig. 2B, C). In contrast, chondrocyte-derived elements (cartilage) were unaffected. As expected, *ezh1^−/−^* larvae had normal osteoblast- and chondrocyte-derived elements (Fig. 2D). *ezh1^−/−^; ezh2^+/+^* and *ezh1^−/−^; ezh2^−/−^* larvae were grossly indistinguishable (Fig. 2C, 2E). Loss of *ezh2* but not *ezh1* significantly reduced BSR3 length, an effect modestly enhanced by combined *ezh1*/*ezh2* deficiency (Fig. 2F). We conclude Ezh2 is the primary zygotically-expressed H3K27me3 methyltransferase during craniofacial dermal bone formation with Ezh1 serving a minor, partially redundant role.

**Figure 2.**
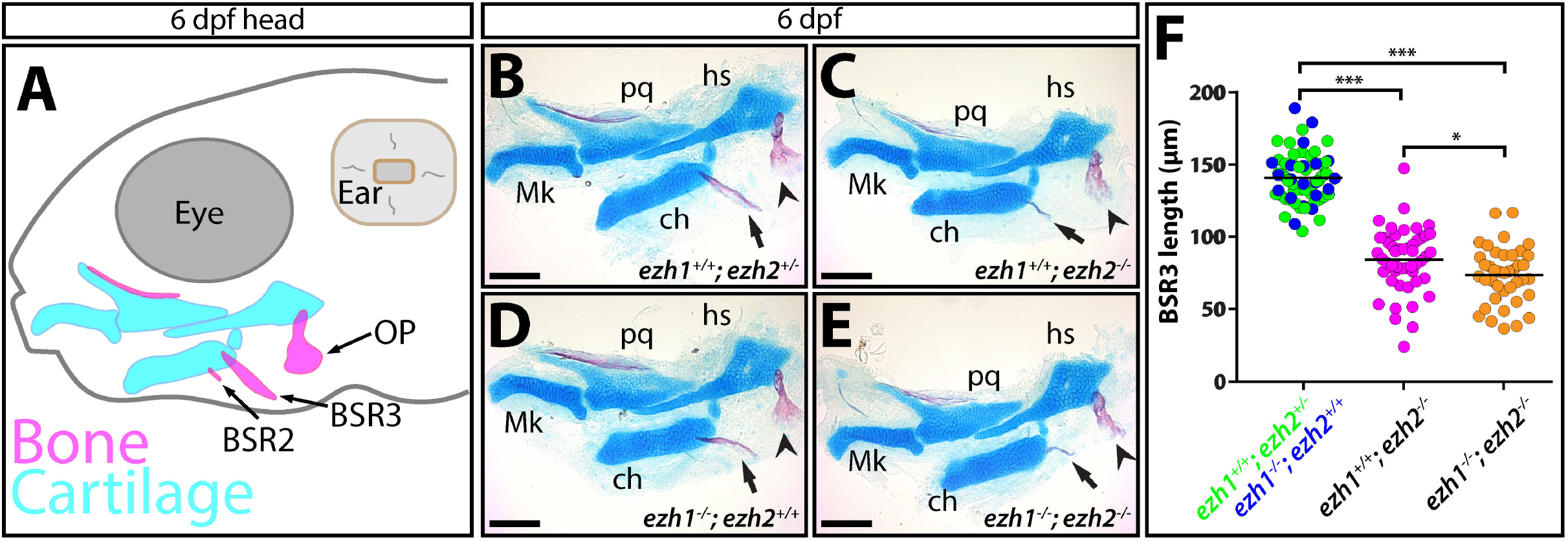
Ezh2 supports craniofacial dermal bone development. **(A)** Schematic of 6 day post fertilization (dpf) zebrafish head. Cartilaginous skeletal elements are in cyan and calcified dermal bone is in magenta. Abbreviations: OP, opercle; BSR, branchiostegal ray. **(B-E)** Flat mount images of alizarin red (red) and alcian blue (blue) stained pharyngeal arch 1 and pharyngeal arch 2 derived skeletal elements from 6 dpf larvae of indicated *ezh1* and *ezh2* genotypes. Abbreviations: Mk, Meckel’s cartilage; pq, palatoquadrate; hs, hyosymplectic; ch, ceratohyal. Arrowhead points to OP; arrow indicates BSR3. Scale bar = 100 μm. **(F)** Scatter plot comparing BSR3 lengths between 6 dpf larvae of the indicated genotype groups. Data points represent individual animals, color-coded by genotype, and means are shown. * = P-value < 0.05, *** = P-value < 0.001 by Student’s two-tailed *t*-tests.

Reduced craniofacial dermal bone in *ezh2^−/−^* larvae could result from incomplete maturation of osteoblasts, a dearth of cranial neural crest cell-derived osteoblast specification and/or migration, or loss of a proliferative pre-osteoblast pool while the bones enlarge. Mouse ES cells and embryonic fibroblasts exhibit reduced osteoblast differentiation in the absence of PRC2 (Chamberlain et al., 2008; Lavarone et al., 2019; Pasini et al., 2007). In addition, Runx2^+^ preosteoblasts of the mouse calvarium fail to fully differentiate in conditional *Ezh2* mutants (Ferguson et al., 2018). We stained 6 dpf *ezh2^−/−^* larvae and wildtype clutchmates carrying the *Tg(sp7:eGFP)* osteoblast reporter (DeLaurier et al., 2010) with alizarin red to assess bone differentiation. *ezh2*-homozygous mutants had a smaller field of *sp7:eGFP*-expressing osteoblasts. However, those osteoblasts remaining produced relatively equivalent amounts of calcified bone as those of *ezh2^+/−^* siblings (Fig. 3A, B). We conclude Ezh2 is not required for terminal bone differentiation per se and likely instead promotes earlier steps in osteogenesis that generate sufficient osteoblast pools.

**Figure 3.**
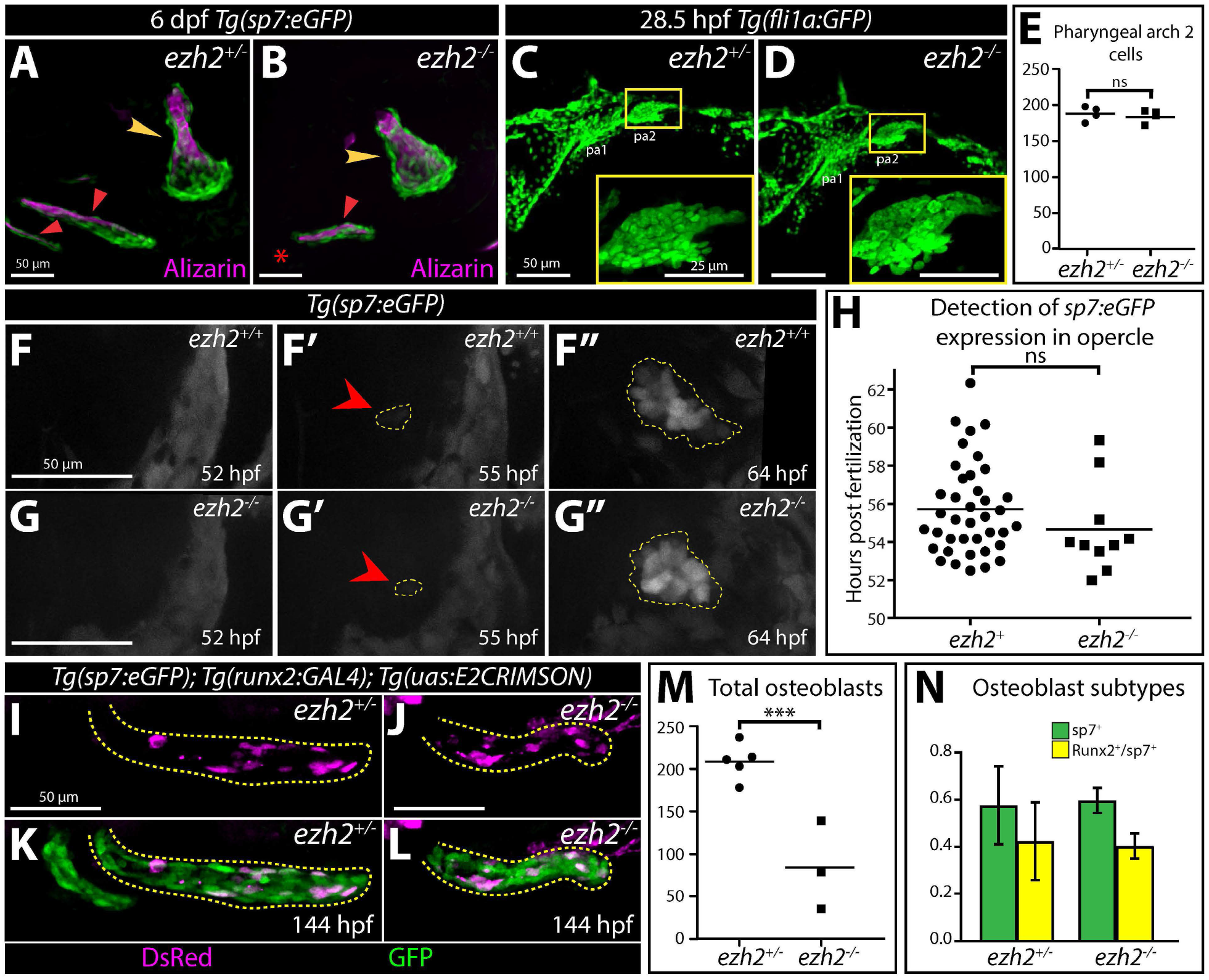
Zygotic Ezh2/PRC2 promotes osteoblast progenitor pool expansion and morphogenesis. **(A, B)** Live confocal images of 6 day post fertilization (dpf) control and *ezh2^−/−^ Tg(sp7:eGFP)* (osteoblasts, green) larvae stained with alizarin red (ossified bone, magenta). Red arrowheads point to branchiostegal rays (BSRs); yellow arrowheads mark opercles (OP); red asterisk highlights the missing BSR2 in the *ezh2^−/−^* larvae. **(C, D)** Confocal images of 28.5 hour post fertilization (hpf) control and *ezh2^−/−^ Tg(fli1a:GFP)* clutchmate embryos immunostained with anti-GFP antibodies marking cranial neural crest cells (green). Inserts show magnifications of pharyngeal arch 2 (pa2), the source of future OP and BSR osteoblasts. **(E)** Scatter plot showing equal number of *fli1a:GFP-labeled* pa2 cells in 28.5 hpf control and *ezh2^−/−^* embryos. **(F-G”)** Still images from time lapse confocal imaging of the developing opercle of wildtype (F-F”) and *ezh2-deficient* (G-G”) *Tg(sp7:eGFP)* embryos (see Supplemental Movie 1). Red arrowheads point to the first GFP^+^ osteoblasts. Osteoblast field outlined in yellow. **(H)** Scatter plot showing unchanged onset of osteoblast maturation, as assayed by *sp7:eGFP* expression, between wildtype and *ezh2* homozygous mutant clutchmates. **(I-L)** Confocal immunofluorescent images of *sp7:eGFP*^+^ (osteoblasts, green) and *runx2:GAL4; uas:E2CRIMSON*^+^ (pre-osteoblasts, magenta) cells of BSR3 (yellow dashed outline) and BSR2 from control and *ezh2*-null 144 hpf clutchmates stained with anti-GFP and anti-DsRed antibodies. **(M)** Scatter plot comparing total number of BSR3 osteoblasts in 144 hpf control and *ezh2^−/−^* larvae. **(N)** Bar graph showing similar proportions of osteoblast subtypes between 144 hpf control and *ezh2^−/−^* clutchmates. Error bars show one standard deviation. Statistical significance in (E, H, M, N) determined by Student’s two-tailed *t*-tests. *** = P-value < 0.005, ns = not significant. Scale bars are 50 μm unless otherwise noted.

We next explored if the deficient dermal bone in the absence of *ezh2* originated from inadequate formation of the precursor progenitor osteoblast field. BSR and opercle osteoblasts are derived from pharyngeal arch 2 (pa2) cranial neural crest cells (CNCCs) (Crump et al., 2006; Crump et al., 2004; Laue et al., 2008). *Ezh2* NCC conditional mutant mice fail to develop craniofacial bones, possibly due to aberrant positional identity precluding CNCC migration (Minoux et al., 2017). Likewise, *Xenopus* CNCCs migrate less upon Ezh2 inhibition (Tien et al., 2015). Using the *Tg(fli1a:gfp)* CNCC marker (Lawson & Weinstein, 2002), we found *ezh2^−/−^* embryos had a normal sized, shaped and CNCC-populated pa2 field at 28.5 dpf (Fig. 3C-E). Therefore, zygotic Ezh2 likely promotes expansion rather than generation of osteoblast-fated pa2 CNCCs.

The net expansion of a progenitor pool depends on proliferation opposed by depletion through differentiation. We evaluated the effects of *ezh2* loss-of-function on pre-osteoblasts (pObs) *in vitro* using the Ezh2-specific small molecule inhibitor EPZ-6438 (Tazemetostat) (Knutson et al., 2013) and AB.9 cells. AB.9 immortalized cells are derived from zebrafish fins (Paw & Zon, 1999) that we newly characterize as Runx2-expressing pObs (Fig. S4A-B). EPZ-6438 effectively inhibited zebrafish Ezh2 as shown by loss of bulk H3K27me3 in treated AB.9 cells (Fig. S4D). Ezh2-inhibited AB.9 cells produced elevated alkaline phosphatase when cultured in differentiation media (Fig. S4E-F), consistent with mammalian cell culture studies (Dudakovic et al., 2016; Dudakovic et al., 2015). Therefore, PRC2 may have a conserved role restraining osteoblast maturation.

Ezh2 opposition of osteoblast maturation *in vitro* anticipates *ezh2^−/−^* embryos would show premature osteoblast differentiation *in vivo.* To test this, we used live time-lapse imaging of *sp7:eGFP* embryos from 52 to 72 hpf to monitor and score the onset of *sp7* expression in forming opercles. Both *ezh2^−/−^* and control clutchmates first showed *sp7*+ cells at the dorsal point of the opercle around 55 hpf (Fig. 3F-H, Supp. Movie 1). The total number of *sp7*-expressing opercle osteoblasts appeared similar through 72 hpf. However, opercle field morphogenesis was abnormal in *ezh2*-deficient animals. Normally, *sp7*+ osteoblasts at the dorsal opercle aspect extend ventrally from two to three dpf before expanding along the anteroposterior axis between 4 and 5 dpf (Brinkley et al., 2016; Eames et al., 2013). In contrast, the opercle field of *ezh2*-null embryos expanded along both axes through 72 hpf (Fig. 3F”, G’’, Supp. Movie 1).

We similarly examined BSR development by imaging *ezh2^−/−^* and control larvae from 76 to 96 hpf (Supp. Movie 2). BSR3-forming osteoblasts initiated *sp7* expression near the ceratohyal and then migrated posteriorly to elongate the field in control animals. In *ezh2^−/−^* larvae, *sp7* expression initiated at the normal time and location but the BSR3 *sp7*+ osteoblast pool largely remained static. Collectively, these live imaging studies argue against abrupt, precocious differentiation of craniofacial bone field osteoblasts in *ezh2* mutants. Further, the abnormal morphogenesis of the bone-forming fields, including deficient BSR3 osteoblast migration, could itself reflect a loss of progenitor cell characteristics. Regardless, we conclude zygotic Ezh2 is dispensable for craniofacial osteoblast differentiation but is required for bone field expansion beyond 72-96 hpf.

Loss of zygotic Ezh2 did not affect pa2 populations at one dpf, initiation of *sp7* expression at two dpf, or the ability of osteoblasts to produce calcified bone. The *sp7*-marked osteoblast field morphogenesis defects seen at two and three dpf likely contributed to shortened bones but do not explain the apparent decrease in total osteoblasts underlying smaller bones that manifests later, between ~4 and 6 dpf. Craniofacial pre-osteoblasts transition from Runx2-to sp7-expressing as they differentiate (Li et al., 2009). We scored *Tg(runx2b:GAL4); Tg(uas:E2CRIMSON)* and *sp7:EGFP-*expressing BSR3 cells at 6 dpf. *ezh2*-null mutants had fewer total osteoblasts but a similar proportion of *runx2/sp7-* and *sp7*-expressing subpopulations (Fig. 3 I-N). We treated *ezh2^−/−^; sp7:eGFP* and clutchmate larvae with EdU from 96 hpf to 98 hpf to assess osteoblast proliferation (Fig. 4A-D). EdU-incorporating osteoblasts were more abundant at the medial to posterior aspect of BSR3 in both control and *ezh2* homozygous mutants. However, *ezh2^−/−^* larvae had half as many EdU-positive osteoblasts (Fig. 4E). We conclude decreased osteoblast proliferation, due to depletion of the pOb pool and/or reduced cell cycling, tempers osteoblast accumulation and causes smaller dermal bones in *ezh2* mutants.

**Figure 4.**
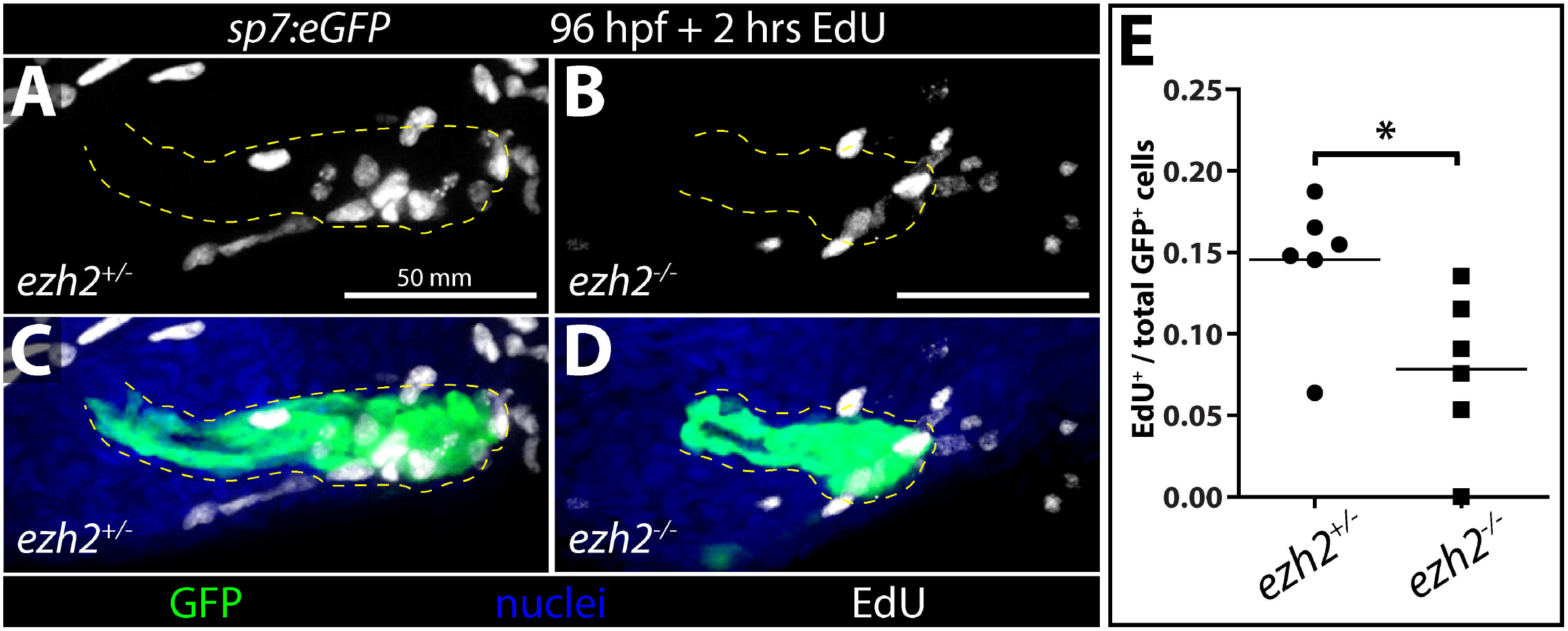
Ezh2 supports craniofacial dermal bone osteoblast proliferation. **(A-D)** Maximum intensity projection confocal images of whole mount 98 hour post fertilization (hpf) control (A and C) and *ezh2*^−/−^ (B and D) *Tg(sp7:eGFP)* larvae stained for EdU incorporation (proliferative, white) and GFP (osteoblasts, green). BSR3 osteoblasts are outlined in yellow. Hoechst-stained nuclei are blue. **(E)** Scatter plot graph showing the fraction of EdU-positive BSR3 osteoblasts of 98 hpf control and *ezh2^−/−^* larvae treated with EdU for 2 hours. P-value is from a two-tailed Student’s *t*-test. * = P-value < 0.05.

### *ezh2^sa1199^* is a hypomorphic allele uncovering vertebrate PRC2 homeotic functions and enabling post-larval stage studies

We obtained the *ezh2^sa1199^* allele and isolated it through extensive outcrossing to rid the background of unknown linked mutations that led to lethality around three dpf, as independently reported (San et al., 2019). An earlier study described *ezh2^sa1199^* as a null allele (Zhong et al., 2018). However, fish homozygous for the *ezh2^sa1199^* allele did not present malformed BSRs or opercles at 6 dpf (Fig. 5A-B). The *ezh2^sa1199^* lesion is a nonsense point mutation at codon 18 that should produce no or non-functional protein. However, we noticed a potential internal translation start site (TSS) at Ezh2 methionine-24 that could generate a truncated protein including the key WD40 binding and catalytic SET domains (Fig. S5A). Indeed, Ezh2 from *ezh2^sa1199/sa1199^* fish was of lower molecular weight and produced at reduced levels (Fig. 5C). Concordantly, *ezh2^sa1199/sa1199^* embryos showed greatly diminished H3K27me3 (Fig. 5C). RT-qPCR showed elevated *ezh2* transcript levels in fish bearing an *ezh2^sa1199^* allele, indicating *ezh2^sa1199^* transcripts were not subject to nonsense mediated decay and rather were stabilized or transcribed at a higher rate (Fig. S5H). *ezh2^sa1199/sa1199^* fish survived to adulthood at near expected ratios (Fig. 5D), albeit with developmental defects including a notable “open mouth” phenotype and fewer bony rays in their anal and pelvic fins (Fig. 5E-G, Fig. S5B-C). Together, we conclude *ezh2^sa1199^* is a hypomorphic allele with substantially reduced protein activity nonetheless capable of supporting near normal development.

**Figure 5.**
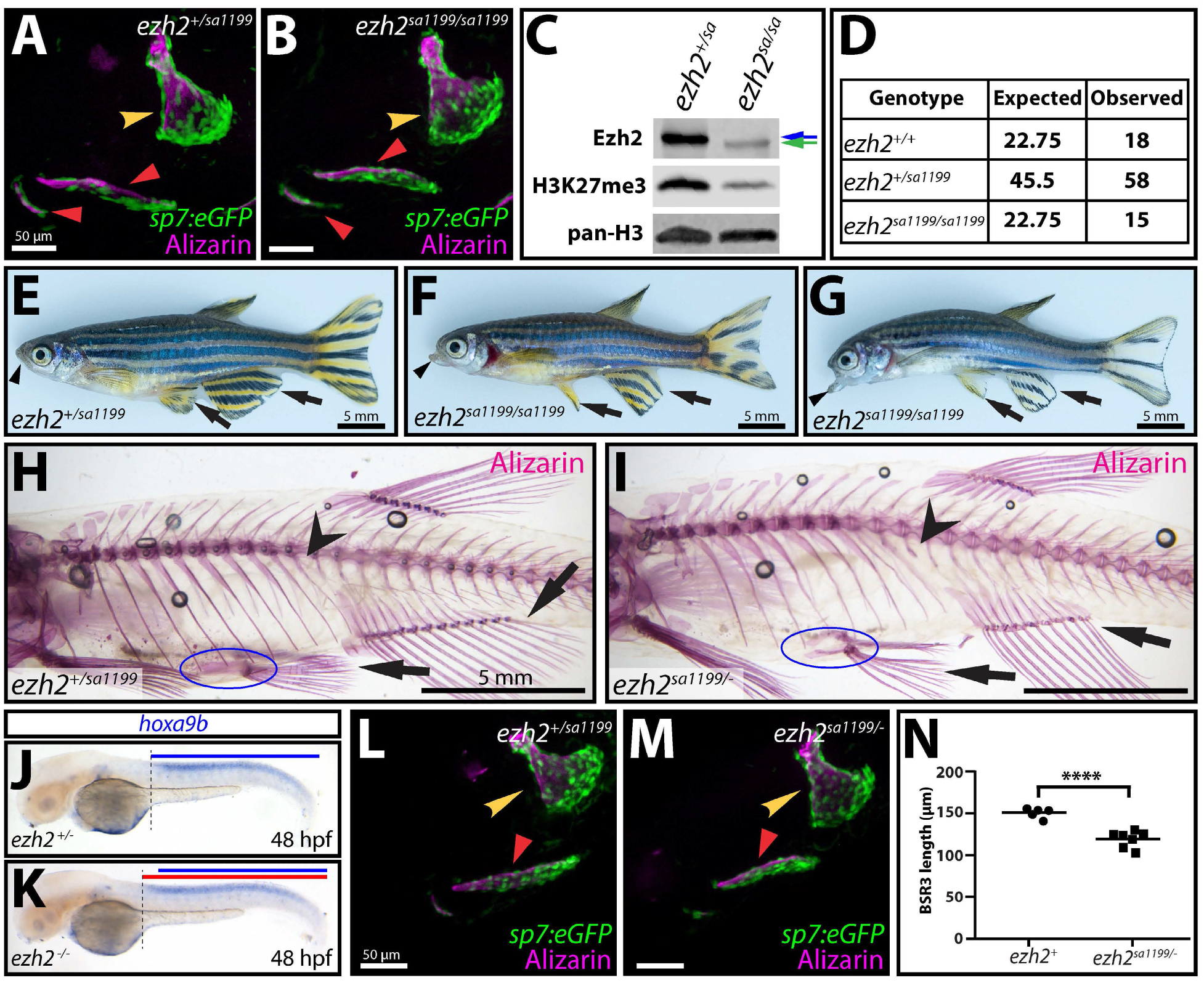
The *ezh2^sa1199^* hypomorph allele unveils appendicular and axial skeletal defects upon PRC2 loss-of-function. **(A, B)** Live confocal imaging of 6 day post fertilization (dpf) wildtype and *ezh2^sa1199/sa1199^ Tg(sp7:eGFP)* (osteoblasts, green) larvae stained with alizarin red (ossified bone, magenta). The branchiostegal rays (red arrowheads) and opercle (yellow arrowhead) are indistinguishable between genotypes. Scale bar = 50 μm. **(C)** Immunoblots of whole cell lysates from 3 day post amputation caudal fin regenerates using Ezh2, H3K27me3, and pan-histone H3 antibodies. Ezh2 protein is reduced and truncated and H3K27me3 lower in *ezh2^sa1199/sa1199^* fins. Blue arrow indicates wildtype Ezh2 and the green arrow points to slightly smaller Ezh2 protein from *ezh2^sa1199^* transcripts. **(D)** Table of expected versus observed number of adult fish of indicated genotypes from an *ezh2^+/sa1199^* in cross. **(E-G)** Whole mount brightfield images of 5 month old wildtype and *ezh2^sa1199/sa1199^* clutchmate fish. Arrows point to anal and pelvic fins. Arrowheads indicate the lower jaw. Scale bar = 5 mm. **(H, I)** Alizarin red-stained skeletal preparations of 4 month old adult (H) *ezh2^+/sa1199^* and (I) *ezh2^sa1199/-^* clutchmate fish. Arrowheads highlight the lost precaudal-identity vertebrae in *ezh2^sa1199/-^* animals. Arrows mark anal and pelvic fins, which have a decreased number of bony rays in *ezh2*-deficient fish. The blue oval circumscribes the reduced pelvic girdle in *ezh2^sa1199/-^* animals. **(J, K)** *hoxa9b* (blue) in situ hybridization of 48 hpf wildtype and *ezh2^−/−^* clutchmate embryos. Blue bar indicates extent of *hoxa9b* in wildtype embryos. Red bar shows the slight anterior expansion of *hoxa9b* in *ezh2* mutant homozygotes. Dotted vertical line marks the anterior *hoxa9b* expression boundary. **(L, M)** Confocal images of live 6 dpf *ezh2^+/sa1199^* and *ezh2^sa1199/-^ Tg(sp7:eGFP)* (osteoblasts, green) larvae stained with alizarin red (ossified bone, magenta). Scale bar = 50 μm. **(N)** Scatterplot showing modest BSR3 length reduction in *ezh2^sa1199/-^* larvae at 6 dpf. **** = P-value < 0.0001 by two-tailed Student’s *t*-test.

While *ezh2^sa1199/sa1199^* adults had mild appendage patterning defects, they lacked overt homeotic transformations seen in other organisms bearing hypomorphic PRC2 mutations. Therefore, we combined our *ezh2* null and *ezh2^sa1199^* hypomorph alleles to look for more severe body pattern defects in *ezh2^sa1199/-^* adults. By alizarin red staining, *ezh2^sa1199/-^* fish had fewer bony rays in their pelvic and anal fins, similar to *ezh2^sa1199/sa1199^* adults, as well as their caudal fins (Fig. S5F-G). 30% of *ezh2^sa1199/-^* fish lacked pelvic fins and pelvic girdles altogether. Further, *ezh2* transheterozygous fish were missing a rib with the most proximal pre-caudal vertebra assuming a more caudal identity (Fig. 5H-I). Similar transformations in vertebrae identity are observed in mice with mutations in PRC1 genes (Akasaka et al., 1996; Coré et al., 1997; Courel et al., 2008; Isono et al., 2005; Katoh-Fukui et al., 1998; Suzuki et al., 2002; van der Lugt et al., 1994) and in an *Eed* hypomorph (Shumacher et al., 1996). However, this is the first homeotic transformation reported in zebrafish upon PcG loss of function. Axial skeleton homeotic phenotypes are often accompanied with anterior misexpression of posterior *hox* genes. Zebrafish *rnf2* (PRC1 subunit) mutants (van der Velden et al., 2012) and *MZezh2^hu567/hu5670^* (Rougeot et al., 2019; San et al., 2016) animals exhibit anterior expansion of the posterior *hox* gene, *hoxa9b.* Accordingly, RNA in situ hybridizations showed modest anterior expansion of *hoxa9b* in *ezh2^−/−^* embryos (Fig. 5J-K).

We reasoned since surviving *ezh2^sa1199/-^* adults had more severe defects than *ezh2^sa1199/sa1199^* fish, *ezh2^sa1199/-^* larvae similarly would present with more severe craniofacial phenotypes. Using alizarin red staining and the *sp7:eGFP* background, we found 6 dpf *ezh2^sa1199/-^* transheterozygotes had shortened BSR3s (Fig. 5L-N). Likewise, *ezh2^sa1199/-^; sp7:eGFP* larvae had slightly smaller opercles with moderately wider opercle necks than wildtype larvae (Fig. 5L,M). Further, *ezh2^sa1199/-^* three week old larvae had diminished cranial calcification (Fig. S5D-E). The additional axial skeleton and fin patterning phenotypes and exacerbated craniofacial defects in adult and larval *ezh2* transheterozygotes further demonstrate *ezh2^sa1199^* is a hypomorph. This allele allows for investigation of post larval roles of Ezh2 and affords a second vertebrate model to study mechanisms of PcG maintenance of segmented Hox gene expression.

### Major Ezh2/PRC2 pleiotropic organogenesis functions are concentrated immediately after maternal/zygotic transition

Maternal zygotic *ezh2* null embryos generated by germline transplantations (*MZezh2*-nulls) have liver, pancreas, brain, eyes, and heart defects, dying around two dpf (San et al., 2016). We considered if *ezh2^sa1199^*, as a hypomorph providing enough function for largely normal development to adulthood, would facilitate studies of combined maternal/zygotic Ezh2 function. We crossed *ezh2^sa1199/sa1199^* females to *ezh2^+/−^* males and reared offspring until 6 dpf (Fig. 6, Fig. S6). Many defects in the *ezh2* maternal zygotic mutants (hereafter referred to as *M^sa1199^Zezh2^sa1199/-^*) corresponded with *MZezh2-null* phenotypes (San et al., 2016). Likewise, we also found zygotic wildtype *ezh2* was sufficient to fully rescue maternal zygotic mutant phenotypes (Fig. 6, Fig. S6). *M^sa1199^Zezh2^sa1199/-^* defects first manifested at two dpf, unlike *MZezh2*-nulls that exhibit phenotypes as early as one dpf (San et al., 2016). *M^sa1199^Zezh2^sa1199/-^* embryos had decreased body length and reduced yolk extension at two dpf. About 25 percent of *M^sa1199^Zezh2^sa1199/-^* embryos had heart edema at 48 hpf, which was fully penetrant by 56 hpf (Fig. S6A-C). 72 hpf *M^sa1199^Zezh2^sa1199/-^* embryos exhibited extreme cardiac edema associated with greatly elongated hearts lacking distinct atrium and ventricle chambers, similar to the “stringy heart” described in *MZezh2*-nulls (San et al., 2016) (Fig. 6C-D). 72 hpf *M^sa1199^Zezh2^sa1199/-^* embryos also had notably small eyes.

**Figure 6.**
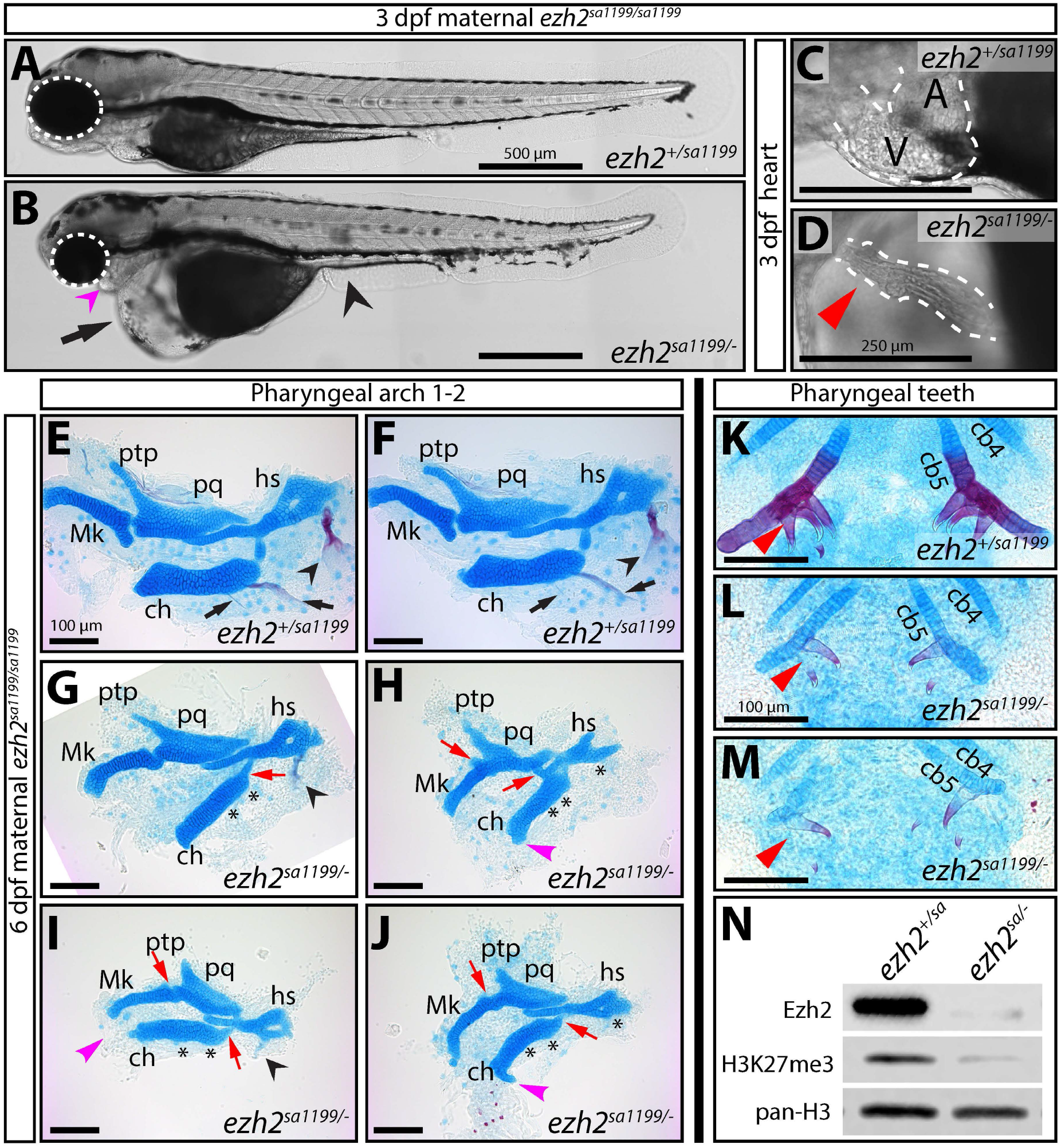
Maternal Ezh2 can fulfill PRC2’s primary developmental roles. **(A, B)** Whole mount differential interference contrast (DIC) images of three day post fertilization (dpf) siblings from a ♀ *ezh2^sa1199/sa1199^* x ♂ *ezh2^+/−^* cross. Zygotic wildtype *ezh2* (A) rescues defects present in maternal-zygotic *ezh2* mutant larvae (B). Arrow points to severe cardiac edema. Black arrowhead indicates absent yolk extension. Magenta arrowhead shows a reduced jaw. White dashed ovals encircle eyes. Scale bar is 500 μm. **(C, D)** Magnified DIC images of hearts from fish in (A) and (B). Dashed white lines outline hearts. (C) *ezh2^+/sa1199^* larvae have a well-defined atrium (A) and ventricle (V). In contrast, (D) maternal *ezh2^sa1199/sa1199^* zygotic *ezh2^sa1199/-^* larvae have extended hearts lacking distinct chambers. Scale bar is 250 μm. **(E-M)** Flat mount preparations showing alizarin red (ossified bone, red) and alcian blue (chondrocyte-derived cartilage, blue) stained skeletal elements from 6 dpf control and maternal-zygotic *ezh2* mutant larvae. Pharyngeal arch 1 and 2 derived elements are normal in zygotic *ezh2* wildtype larvae (E, F). Maternal *ezh2^sa1199/sa1199^* zygotic *ezh2^sa1199/-^* clutchmates (G-J) have fusions of cartilaginous elements and lack nearly all ossification. Black arrows point to branchiostegal rays. Black arrowheads indicate opercle bones. Asterisks denote lost osteoblast-derived elements in maternal-zygotic *ezh2* mutants. Red arrows point to fusions between chondrocyte-derived elements. Magenta arrowheads mark ectopic growth of the anterior process of the ceratohyal. Abbreviations: Mk, Meckel’s cartilage; pq, palatoquadrate; ptp, pterygoid process; hs, hyosymplectic; ch, ceratohyal. Scale bar is 100 μm. **(K-M)** Skeletal preparations showing ceratobranchial 4 and 5 and pharyngeal teeth of control and maternal-zygotic *ezh2* mutant clutchmates. Red arrowheads indicate pharyngeal teeth. Abbreviations: cb, ceratobranchial. Scale bar is 100 μm. **(N)** Whole cell lysate immunoblots from pooled three dpf control and maternal zygotic *ezh2* mutant larvae using Ezh2, H3K27me3, and pan-histone H3 antibodies.

*M^sa1199^Zezh2^sa1199/-^* four to six dpf larvae were exceptionally edematous and exhibited additional organogenesis defects. The intestinal bulb was absent and there was no distinct swim bladder (Fig. S6D-G). *M^sa1199^Zezh2^sa1199/-^* intestinal cells appeared compact instead of adopting a long columnar shape and organization. The liver was elongated and lacked its bilateral configuration (Fig. S6I-J). The pancreas appeared thin and did not establish its characteristic “delta shape” at the anterior end (Fig. S6H, J). Skeletal muscle fibers were wavy, a phenotype not described in *MZezh2*-nulls presumably because they die before the defect could be observed (Fig. S6E, G).

*M^sa1199^Zezh2^sa1199/-^* embryos and larvae largely failed to elongate their jaws, indicating more severe craniofacial defects than observed in *ezh2^−/−^* larvae (Fig. 6A-B, Fig. S6D-G). Flat mount skeletal preparations confirmed 6 dpf *M^sa1199^Zezh2^sa1199/+^* larvae had no defects in pa1/2-derived elements (Fig. 6E-F). In contrast, *M^sa1199^Zezh2^sa1199/-^* clutchmates lacked BSRs and their opercles were missing or diminished (Fig. 6G-J). Osteoblast-derived pharyngeal teeth were also underdeveloped (Fig. 6K-M, Fig. S7). *M^sa1199^Zezh2^sa1199/-^* fish also had multiple cartilage defects. Mandibular and hyoid cartilaginous elements were small and commonly fused (Fig. 6G-J). The pterygoid process of the palatoquadrate did not extend, giving it a short stubby appearance. The ceratohyal was smaller and had abnormal cartilage growth at the ventro-anterior tip where the ventral hypohyal bone typically develops (Fig. 6G-J). Further, the anterior end of the basihyal did not adopt its typical fan-like shape (Fig. S7). We earlier described how smaller and misshapen dermal bones upon loss of zygotic *ezh2* are likely caused by the progressive depletion of progenitor osteoblasts. The considerable cartilage and profound calcified bone skeletal defects in *M^sa1199^Zezh2^sa1199/-^* embryos suggests Ezh2/PRC2 has an earlier function affecting development of pa1/2-associated CNCC cells and/or development of both osteoblast and chondrocyte lineages.

*M^sa1199^Zezh2^sa1199^* embryos had similar pleiotropic organogenesis defects but less and more severe than those of *MZezh2^hu5670/hu5670^* embryos and *ezh2^−/−^* larvae, respectively. We detected very low levels of Ezh2 in whole cell lysates of 3 dpf *M^sa1199^Zezh2^sa1199/-^* larvae (Fig. 6N). Correspondingly, there was a substantial decrease in bulk H3K27me3 (Fig. 6N). Residual Ezh2^sa1199^ protein and activity likely led to less severe defects of *M^sa1199^Zezh2^sa1199/-^* larvae compared to *MZezh2*-nulls. In comparison, even though Ezh2 was undetectable and H3K27me3 greatly decreased, 6 dpf zygotic *ezh2*^−/−^ larvae had only mild defects (Fig. 1K, M). This contrast along with the absence of phenotypes in *M^sa1199^Zezh2^+/sa1199^* larvae (Fig. 6, Fig. S6), which seemingly have almost no maternal Ezh2 activity, highlights maternal *or* zygotic Ezh2 can fulfill PRC2’s most critical developmental functions. Further, PRC2 activity is most essential immediately after zygotic genome activation when both maternal and zygotic Ezh2 pools are present.

The residual H3K27me3 in 3 dpf *M^sa1199^Zezh2^sa1199/-^* embryos detected by Western blot could reflect remaining activity of the truncated Ezh2^SA1199^ protein or Ezh1 manifesting in variable changes in bulk H3K27me3 in different cell types. We immunostained 3 dpf *M^sa1199^Zezh2^sa1199/-^* embryos and zygotic wildtype clutchmates with antibodies against H3K27me3 as well as MF20 (myosin heavy chain) and pan-cadherin to accentuate tissue organization. We examined gut, eye, skeletal muscle, spinal cord, chondrocytes and heart (Fig. S8). In all cases, bulk H3K27me3 was undetectable in *M^sa1199^Zezh2^sa1199/-^* embryos. Combined with our genetic studies, this result indicates: 1) the *ezh2^sa1199^* hypomorph produces insufficient PRC2 activity to sustain detectable, bulk H3K27me3 but nevertheless is considerably less deleterious than a *ezh2* null allele, 2) Ezh1 provides little H3K27me3 activity up to 3 dpf and is unable to even minimally replace zygotic Ezh2 when maternal Ezh2 is unavailable to initiate H3K27me3 marks, 3) PRC2/H3K27me3 is dispensable for body plan establishment and initial organ specification whereas at least some PRC2 activity and H3K27me3 broadly supports organ expansion, cell diversification, and morphogenesis, and 4) minimal Ezh2/PRC2 and H3K27me3 are sufficient for near normal development to adulthood, provided Ezh2/PRC2 can establish initial H3K27me3 marks upon zygotic genome activation.

## DISCUSSION

We explore genetic contributions of zebrafish Ezh1 and Ezh2 H3K27me3 methyltransferases to reveal core vertebrate PRC2 roles and strengthen the framework for zebrafish PcG research. We show Ezh2 provides major PRC2-activities during embryogenesis with only minor, redundant and non-compensatory contributions by non-maternally deposited Ezh1. Indeed, Ezh1 is not required for viability or fertility despite progressively supplanting Ezh2 in relative expression after embryogenesis. Either maternal or zygotic Ezh2 can provide early embryonic PRC2 function, showing the importance of establishing H3K27me3 landscapes immediately after zygotic genome activation. Nonetheless, PRC2 is dispensable for gastrulation and body plan establishment. We show how abnormally sized and shaped craniofacial dermal bones in *ezh2* zygotic nulls arise from reduced proliferation and disrupted migration of progenitor osteoblasts, which otherwise are specified normally and can differentiate into bone-producing cells. An *ezh2* allelic series reveals additional Ezh2-PRC2 organogenesis roles and uncovers axial skeletal homeotic transformations upon zebrafish PcG disruption. Further, after the early embryonic critical period, little PRC2 activity – unable to sustain bulk H3K27me3 – is sufficient for much of development to adulthood. Collectively, our study suggests broad but ancillary maintenance rather than instructive roles for PcG-mediated repression in cell fate determination.

### Ezh1 is dispensable for zebrafish development and cannot substitute for Ezh2/PRC2

We confirm reports *ezh2* mRNA is maternally deposited and highly expressed throughout the first five days of zebrafish development (Chrispijn et al., 2018; San et al., 2018; San et al., 2016; Sun et al., 2008; White et al., 2017). *ezh2* expression then progressively decreases through larval development. Contrasting one report (Völkel et al., 2019), we and others found *ezh1* mRNA is not maternally contributed (Chrispijn et al., 2018; San et al., 2018; San et al., 2016; Sun et al., 2008; White et al., 2017). Zygotic *ezh1* expression steadily increases from one to five dpf, exceeding *ezh2* expression by four dpf (Chrispijn et al., 2018; San et al., 2018; San et al., 2016; Sun et al., 2008; White et al., 2017). Such complementary expression profiles are consistent with Ezh2 serving as the primary H3K27me3 methyltransferase in progenitor and highly proliferative cells whereas Ezh1 typically assumes the H3K27me3 maintenance role in mature cell populations (Ezhkova et al., 2009; Margueron et al., 2008).

While *ezh2^−/−^* fish die around two weeks post fertilization, *ezh1^−/−^* animals are viable and fertile, matching mouse and previous zebrafish studies (Ezhkova et al., 2011; Völkel et al., 2019). Ezh1 therefore does not have notable, unique developmental roles. Ezh2 may substitute for Ezh1 at later life stages despite its relatively lower expression. In contrast, embryonic lethality of *ezh2* single homozygous mutants shows that Ezh1 cannot fully accommodate PRC2 functions (San et al., 2016). We now show that *ezh1; ezh2* double mutants have nearly identical phenotypes as *ezh2* single mutants. This could reflect distinct *ezh1* and *ezh2* temporal expression patterns. *ezh2* always co-expresses with *ezh1* while *ezh1* only coincides with *ezh2* after one dpf. Alternatively, Ezh1/PRC2 and Ezh2/PRC2 may have distinct biochemical activities (Margueron et al., 2008). Ezh1 and Ezh2 are equally capable of depositing H3K27me1 (Lavarone et al., 2019; Lee et al., 2018). However, Ezh2 more readily catalyzes H3K27me2/3 methylation, especially in the presence of cofactors and allosteric activators (Lavarone et al., 2019; Lee et al., 2018; Lee et al., et al., 2018). Further, Ezh2/PRC2 complexes may have unique ability to catalyze spreading of H3K27me3 marks from nucleating centers (Lavarone et al., 2019). Our observation Ezh1 can partially replace Ezh2/PRC2-deposited bulk H3K27me3 marks in zygotic but not maternal/zygotic *ezh2* mutants further indicates Ezh1/PRC2 may be able to maintain but not initiate H3K27me3 marks.

### *ezh2^sa1199^* is a useful hypomorph for uncovering PRC2 organogenesis roles

We correct misconceptions about the *ezh2^sa1199^* allele by showing how its predicted nonsense and null lesion instead produces a severe hypomorph due to faint internal translation initiation. Zhong and colleagues define and use *ezh2^sa1199^* as a null to investigate hematopoiesis and clock gene expression (Zhong et al., 2018). In complete contrast, San et al. conclude the *ezh2^sa1199^* allele is not a loss of function and further state this allele produces no phenotype (San et al., 2019). Zhong et al. assumed the *ezh2^sa1199^* allele is a null based on immunoblots. However, their studies present phenotypes likely caused by an uncharacterized linked mutation in the Sanger Project line. San et al., like us, isolated *ezh2^sa199^* away from the linked mutation (San et al., 2019). In contrast to their findings, we show *ezh2^sa1199^*, while not a null, produces little Ezh2 protein and activity.

We show how *ezh2^sa1199^* produces minimal but functional protein in spite of a nonsense mutation at codon 18 by weak internal translational initiation upstream of Ezh2’s essential WD40 binding and catalytic SET domains. The minute amount of truncated Ezh2 produced by *ezh2^sa1199^* is sufficient to fulfill much of Ezh2/PRC2’s embryonic roles. Therefore, the *ezh2^sa1199^* allele exemplifies how translational machinery can bypass early termination signals to convert expected null alleles into hypomorphs without severe phenotypes. Relatedly, the allele provides a design framework to intentionally produce valuable hypomorphs.

*ezh2^sa1199/-^* zebrafish can develop to adulthood even with almost complete loss of zygotic Ezh2/PRC2 activity. Therefore, core vertebrate PRC2’s transcriptional maintenance roles seem surprisingly superfluous, at least in a laboratory setting, once past a critical early embryonic window. Further, the relatively minor phenotypes of *ezh2^sa1199/-^* adults and *ezh1^−/−^; ezh2^−/−^* larvae contrasted by major bulk H3K27me3 deficiency imply sparse H3K27me3 promoter-centric nucleation versus H3K27me2/3 spreading mostly determines gene repression, as seen in ES cells (Lavarone et al., 2019). Interestingly, Lavarone et al. propose ES cell differentiation defects in the absence of H3K27me are due to the loss of broad H3K27me2 domains and resultant accumulation of H3K27 acetylation rather than derepression. While we have not assessed H3K27me2 or H3K27ac, the generalized disruption of zebrafish embryogenesis in *ezh2* mutants could reflect a similar secondary effect meriting further investigation.

*ezh2^sa1199^* homozygous females are fertile, facilitating studies of maternal Ezh2 during embryogenesis. When crossed to *ezh2^+/−^* males, offspring exhibit phenotypes similar to maternal-zygotic *ezh2* mutants described in previous work (San et al., 2016), albeit with a range of expressivity and surviving up to 6 dpf. Maternal *ezh2^sa1199/sa1199^* zygotic *ezh2^sa1199/-^* mutants enabled us to evaluate organ development at later time points than complete maternal-zygotic *ezh2*-nulls (San et al., 2016). All together, *ezh2^sa1199^* is a severe and informative hypomorph.

### PRC2 has conserved roles in craniofacial, axial, and appendicular skeletal patterning

Our study of craniofacial skeletal abnormalities in *ezh2^−/−^* mutants uncovers how PRC2 enables expansion of dermal bone pre-osteoblast pools and their morphogenesis into correctly sized and shaped bones. *ezh1^−/−^; ezh2^−/−^* larvae develop slightly smaller dermal bones, supporting only minimal Ezh1/Ezh2 redundancy for this PRC2 function. The small craniofacial dermal bone phenotypes correlate with a decrease in the fraction of actively dividing pObs. However, PRC2 seems unlikely to directly promote cell cycle entry given the essential window for PRC2 function precedes the onset of overt proliferation and bone defects. Dermal bone defects worsen and craniofacial cartilage bone phenotypes manifest in *M^sa1199^Zezh2^sa1199/-^* mutants, suggesting a shared PRC2 role in cartilage/bone cells early in embryogenesis. We also found enhanced osteoblast maturation of pObs in vitro upon Ezh2 inhibition. Together, we suggest Ezh2/PRC2 maintains osteoblast progenitor characteristics, consistent with links of PRC2 with progenitor state maintenance (reviewed in Scelfo et al., 2015).

Mouse conditional knockout models disrupting PRC2 develop reduced craniofacial skeletal elements reminiscent to phenotypes we describe in *ezh2^−/−^* zebrafish. Conditional ablation of *Ezh2* in cranial neural crest cells using *Wnt1:Cre* causes severe craniofacial defects, including reduced calcification, a poorly formed styloid process, and a missing hyoid bone (Schwarz et al., 2014). Both are BA2/PA2 derived bones like fish BSRs and OPs. In addition, *Pdgfra:CreER-mediated* deletion of *Ezh2* in cranial mesenchymal stem cells decreases calvarial bone calcified matrix (Ferguson et al., 2018). In this mammalian context, osteoblast progenitors form but fail to fully differentiate, unlike the smaller osteoblast pools with decreased proliferation and retained differentiation we describe for *ezh2^−/−^* zebrafish craniofacial dermal bone. Therefore, common Ezh2/PRC2 roles in cranial neural crest cell-derived skeletal development may be coincidental rather than reflecting the conserved regulation of specific cell behaviors.

The PcG famously represses *Hox* genes to maintain Hox codes conferring body segmental identities. Many PRC1 mouse mutants have been characterized with axial skeleton homeotic transformations. Embryonic lethality of homozygous null alleles of the three core PRC2 subunits (Eed, Suz12, Ezh2) precludes straightforward such observations for PRC2. Only an *Eed* hypomorph has been reported to cause posterior transformations of the axial skeleton (Shumacher et al., 1996). Our viable hypomorph *ezh2^sa/-^* trans-heterozygote zebrafish enable investigating how PRC2 contributes to axial skeletal and appendage development in another vertebrate. Similar to the mouse *Eed* hypomorph, our zebrafish *ezh2* hypomorphs have posterior transformations with the posterior pre-caudal vertebrae assuming a caudal identity characterized by rib loss. Correspondingly, mutations in *Jmjd3/KDM6A,* an H3K27me3 demethylase, cause anterior homeotic transformations and gain a rib (Naruse et al., 2017). We also see changes in appendage morphology, and in some cases, entirely missing pelvic fins. Numerous links of *Hox* genes to appendage patterning (Jain et al., 2018; Minguillon et al., 2012; Nakamura et al., 2016) suggest misexpression of *hox* genes could account for these *ezh2* loss-of-function phenotypes.

### Ezh2 contributes essential PRC2/H3K27me3 function concentrated post-genome activation and not reflected by bulk H3K27me3

San et al. showed *MZezh2*-null embryos form a regular body plan and many organs retain early specification markers (San et al., 2016). Our genetic studies combining *ezh1* and *ezh2* loss confirm the *MZezh2*-null phenotype represents the total loss of PRC2 from oocyte through embryonic lethality at 2 dpf. This genetic approach neatly complements a recent multi-omics study showing near loss of H3K27me3 in absence of maternal and zygotic Ezh2 but with retained Ezh1 (Rougeot et al., 2019). The surprisingly normal body plan of *MZezh2* mutant embryos indicates PRC2 does not play central, instructive roles in earliest steps of embryogenesis including initial cell lineage specification or gastrulation given presence of organs from all germ layers. Nevertheless, PRC2’s key organogenesis functions seem concentrated during a narrow window of development succeeding zygotic genome activation. Either maternal or zygotic Ezh2 can support these PRC2 functions, underscored by the wildtype phenotype of *M^sa1199^Zezh2^+/sa1199^* larvae compared to severe defects of *M^sa1199^Zezh2^sa1199/-^* clutchmates. Consistently, ChIP-Seq studies find only low levels of H3K27me3 prior to zygotic genome activation (Lindeman et al., 2010; Murphy et al., 2018; Vastenhouw et al., 2010; Zhang et al., 2014).

Relatively normal larval development and adult homeostasis with profound H3K27me3 loss alternatively could suggest H3K27me3 is not required for continuous repression of most H3K27me3-marked genes. Curiously, this appears the case for ancestral PRC2/H3K27me3 studied in *Neurospora* (Jamieson et al., 2013) and, as previously noted, ES cells (Lavarone et al., 2019). H3K27me3 marks then may represent fallback barriers against rare, spurious gene activation still largely dependent on collective transcription factor activity. This concept could explain why H3K27me3 establishment is especially important early in development when extensive gene regulatory events simultaneously promote an array of cell behaviors, including multifarious cell specification. Further, this model portends a supporting, perhaps non-specific, role for H3K27me3 *deposition* in gene repression while allowing for instructive H3K27me3 *removal* for gene activation switches. Exemplary zebrafish studies link Kdm6b (Utx/Jmjd3) H3K27me3 demethylases to cardiomyocyte specialization during heart ventricle development (Akerberg et al., 2017) and initiation of fin regeneration (Stewart et al., 2009). Regardless, varied in vivo PRC2 organogenesis roles then represent a passive, generalized PRC2 role maintaining cell states – e.g. progenitor vs. differentiated – as well as cell and positional identities. PRC2 loss then manifests in differential phenotypes reflecting the particular gene regulatory networks, if any, subject to transcriptional noise in a given context.

## MATERIALS AND METHODS

### Zebrafish

The University of Oregon Institutional Animal Care and Use Committee (IACUC) approved and monitored all zebrafish procedures following the guidelines and recommendations in the Guide for the Care and Use of Laboratory Animals (National Academic Press). The following established lines were used in this study: *Tg(sp7:EGFP)b1212* (DeLaurier et al., 2010), *Tg(runx2b:GAL4)* (gift from Chuck Kimmel), *Tg(uas:E2CRIMSON)b1229* (Nichols, Pan, Moens, & Kimmel, 2013), *Tg(fli1a:EGFP)y1* (Lawson & Weinstein, 2002), and *ezh2^sa1199^* (Kettleborough et al., 2013). *ezh2^b1392^* and *ezh1^b1394^* were newly generated as described below.

### CRISPR/Cas9 generation of mutant alleles

*ezh2^b1392^* and *ezh1^b1394^* were generated by CRISPR/Cas9-mediated targeting following procedures previously detailed (Akerberg et al., 2017). Briefly, Cas9 mRNA and guide RNAs (gRNAs) were microinjected into one-cell stage embryos. F0 animals were outcrossed to recover germline-targeted progeny identified by amplicon sequencing. Stable lines were established by repeated outcrossing. gRNA targeting sequences were: 5’-AATGGACTGGATCCGGAGCT-3’ for *ezh1* and 5’-GTTCCACGCAAGGAGCTCAC-3’ for *ezh2*.

### Genotyping

Fish were genotyped by PCR using primers flanking the mutated region followed by restriction enzyme digests that distinguish between wildtype and mutant amplicons. CRISPR/Cas9-generated indels disrupted BspEI and SacI restriction sites in *ezh1* and *ezh2* mutant alleles, respectively. CRISPR mutant genotyping primers: *ezh1* 5’-GGCTGAATGTTTGTGGCTTTTTCACC-3’, 5’-GGAAATGGCGAGGCAAAGCTGAC-3’, *ezh2* 5’-GTAATCCATGGGCTAGTACAGC-3’, 5’-CCCTTTCACTTTTAACACTGTGC-3’. Primers to genotype *ezh2^sa1199^* introduced a BstUI restriction site in amplicons of the wildtype allele (base in bold and underlined within the reverse primer). Primers: *ezh2^sa1199^* 5’-CATGGACATCTTTGGGTCCT-3’, 5’-AGCCGCATGTACTCAGACTTCAC**<u>G</u>**C-3’

### Skeletal staining

Alcian Blue and Alizarin Red staining of zebrafish larvae was performed as described (Walker & Kimmel, 2007) with subsequent imaging using a Leica DM4000B upright microscope (flat mount).

For live Alizarin Red staining, 6 dpf fish were placed in embryo medium (EM) containing 0.005% Alizarin Red and 10 mM HEPES for 1 hour in the dark. Fish were rinsed for 15 min in EM, anesthetized with tricaine, mounted in 1% low melt agarose containing tricaine, and imaged using a Leica SD6000 spinning disk confocal microscope with a 20x objective.

Juvenile and adult fish 3 weeks to 7 months old were euthanized with tricaine until opercle movement ceased for at least 10 minutes and then fixed in 2% PFA/PBS for 4 hours to overnight at room temperature. Fish were washed for 1 hour with 100 mM Tris pH 7.5 / 25 mM MgCl_2_ and then incubated overnight in 0.02% Alcian Blue / 80% ethanol / 100 mM Tris pH 7.5 / 25 mM MgCl_2_. Samples were washed sequentially with 80% ethanol/ 100 mM Tris pH 7.5 / 25 mM MgCl_2_,50% ethanol / 100 mM Tris pH 7.5, and 25% ethanol / 100 mM Tris pH 7.5 for one hour each. 3% H_2_O_2_ / 0.5% KOH was used to bleach samples until eyes became light brown. Samples were washed with 35% saturated borate for one hour, and then cleared with 1% trypsin / 35% sodium borate. Cleared samples were washed for one hour with 10% glycerol / 0.5% KOH and stained with 0.02% alizarin red/ 10% glycerol / 0.5% KOH overnight. The next day, samples were washed with 50% glycerol / 0.5% KOH for one hour. Samples were then washed overnight with fresh 50% glycerol/ 0.5% KOH. Skeletons were imaged using a Leica M165 FC stereomicroscope.

### Immunoblotting

Three and 6 dpf larvae were bisected using the vent as a landmark. Anterior portions were stored in 15 μL RIPA buffer at −80°C. Posterior ends were placed in 30 μl 50 mM NaOH and boiled at 95°C for 20 minutes to process for genomic DNA. These samples were neutralized with 7.5 μl 1 M Tris pH 7.5 and then used for genotyping. RIPA-stored samples were pooled according to genotype and lysed using a Bioruptor on high for 5-10 minutes with 30 sec ON/OFF cycles. Individual fin regenerate samples were collected at 72 hours post amputation and placed in 40 μl RIPA buffer and dissociated by pipetting. Once fin tissue was uniformly disrupted, the samples were placed in a Bioruptor for 5 minutes on high for 5 minutes with 30 sec ON/OFF cycles. Samples were centrifuged at 14,000 RPM for 5 minutes. Supernatants were pipetted to fresh tubes with an aliquot reserved to quantify protein concentration by Bradford assay.

Samples were boiled for 10 min at 95°C with 3 μL of 6x loading buffer (300 mM Tris pH 6.8, 60% glycerol, 12% SDS, 0.6M DTT, 0.025% Bromophenol blue), iced for 5 min, and loaded on either a 10% or 15% Tris/Glycine gel in running buffer (25 mM Tris, 144 mM Glycine, 0.1% SDS). Protein was transferred to PVDF membrane in transfer buffer (25 mM Tris, 192 mM Glycine, 20% Methanol) for 2-3 hours. Membranes were rinsed in Tris-buffered saline containing 0.1% Tween-20 (TBST) and blocked with either 5% Bovine Serum Albumin (BSA), or 10 % dry milk in TBST for at least 1 hour. Blocked membranes were incubated overnight at 4°C with primary antibodies diluted in blocking solution: rabbit anti-Ezh2 (Cell Signaling Technologies; 1:1000), rabbit anti-H3K27me3 (Millipore, 07-449, Lot #2064519; 1:2000), and rabbit anti-histone H3 (Abcam; 1:10000). The next day, membranes were rinsed with TBST, washed four times with TBST for 10 minutes each, and then incubated with HRP-conjugated secondary antibodies for two hours at room temperature. Wash steps were repeated. Immunoblots were developed with ECL (GE) and imaged using a Li-COR Odyssey.

### Time-lapse imaging

Offspring from an *ezh2^+/−^; Tg(sp7:eGFP)* x *ezh2^+/−^* cross were anesthetized with 80 mg/l clove oil in embryo medium (EM) and mounted in 0.5% low melt agarose within a glass bottom chamber. Mounted fish were then bathed with EM / clove oil. Fish were imaged every 20 min from 52 hpf to 3 dpf on a Leica SD6000 spinning disk confocal microscope outfitted with a heated stage set to 30°C.

### Quantitative reverse transcription polymerase chain reaction (RT-qPCR)

cDNA was synthesized using Maxima H Minus Reverse Transcriptase (Thermo Fisher) with Trizol-isolated RNA from pooled samples of three genotyped 48 hpf embryos, 6 dpf larvae, or individual 72 hpa fin regenerates. RT-qPCR was performed using KAPA SYBR FAST qPCR master mix reagents (Kapa Biosystems). Relative mRNA expression was normalized using *rpl8* to calculate ΔCTs (threshold cycles). Transcript levels were compared using a ΔΔCT approach. ΔCT values and Student’s two-tailed *t*-tests were used to determine significance. qPCR primer sequences: *rpl8* 5’-CCGAGACCAAGAAATCCAGA-3’, 5’-GAGGCCAGCAGTTTCTCTTG-3’, e*zh2* 5’-CTTGCGTGGAACCAGAGAG-3’, 5’-GCATTCACCAACTCCACAAA-3’, *ezh1* 5’-GAGCTGCTGAATGAAGAATGGTCC-3’, 5’-CCTCCACCATGAAGTTCTGCTG-3’.

### 5-ethynl-2-deoxyuridine labeling

Fish were incubated with 0.1 mg/ml of 5-ethynl-2-deoxyuridine (Thermo Fisher) in embryo media for 2 hours prior to harvesting at 98 hpf. The embryos were fixed with 4% PFA in PBS overnight at 4°C and then washed 3 x 10 minutes in PBS. Embryos then were dehydrated through a methanol series and stored at −20°C for at least 24 hours in 100% methanol. Embryos were rehydrated into PBS containing 0.1% Tween-20 prior to performing EdU Click-It reactions (Thermo Fisher) as described by the manufacturer. Immunostaining was carried out after completing the Click-It reaction. EdU-positive BSR3 osteoblasts, identified by *Tg(sp7:EGFP),* were manually located and counted through image stacks of individual larvae. EdU/GFP double positive cells were normalized to total GFP-positive BSR3 cells and significance was determined by Student’s two-tailed *t*-test.

### Whole mount immunofluorescence

Embryos and larvae were euthanized with tricaine and placed on ice for 10 minutes, fixed overnight at 4°C with 4% PFA in PBS containing 0.1% Tween-20 (PBST), and then washed 3 x 10 minutes with PBST. Samples were dehydrated through a methanol series into 100% methanol and stored at −20°C for at least 24 hours. Samples were rehydrated into PBST and then treated with 10 μg/ml proteinase K diluted in PBST for 30 minutes at room temperature. Samples then were fixed for 20 minutes at room temperature with 4% in PBST, washed 4 x 10 minutes with PBST, and blocked for two hours in 10% normal goat serum/ 1% bovine serum albumin / 1% DMSO / 1X PBS / 0.5% Triton X-100. Samples were incubated with chicken anti-GFP 1:1000 (Aves) and rabbit anti-DsRed 1:500 (Clontech/Takara) antibodies diluted in blocking solution overnight at 4°C. After primary antibody incubations, samples were washed 4 x 15 minutes in PBS containing 0.5% Triton X-100 (PBSTx). Alexa-conjugated secondary antibodies were diluted in blocking solution 1:1000 and incubated with samples for two hours at room temperature. Samples were washed two times for 15 minutes with PBSTx and then incubated with TO-PRO-3 (Thermo Fisher) diluted 1:2000 in PBS for 30 minutes. After three 10 minute PBS washes, samples were mounted in 1% low-melt agarose and imaged using a Leica SD6000 spinning disk confocal microscope.

### Section histology and immunostaining

Embryos and larvae were euthanized at desired timepoints and fixed with 4% PFA in PBST for either 3 hours at room temperature or overnight at 4°C. Samples were washed 3 x 10 minutes with PBST and then dehydrated into 100% ethanol. Samples remained in 100% ethanol overnight, were washed twice in xylenes for 10 minutes, and then embedded in paraffin. Samples were sectioned at 7 μm thickness. Hematoxylin and eosin (H+E) staining followed standard procedures (Ricca Chemical Company). Images were acquired using a Leica DM4000B upright microscope. For immunofluorescent staining, sections were de-paraffinized by washing in xylenes 2 times for 10 minutes. Sections were then rehydrated into MilliQ water and washed with PBST for 5 minutes. Antigen retrieval was performed using 1 mM EDTA and 0.1% Tween-20 and pressurized heating for 5 minutes. Samples were then rinsed three times with PBST and blocked with 10% non-fat dry milk in PBST for one hour at room temperature. Samples were incubated with primary antibodies diluted in blocking solution overnight at 4°C. Primary antibodies were: H3K27me3 (Cell Signaling Technologies, clone C36B11; 1:1000); myosin heavy chain (MF20, Developmental Studies Hybridoma Bank; 1:250); pan-Cadherin (Novus; 1:250). Slides were then rinsed 2 times followed by three 5 minute washes in PBST. Sections were incubated in 0.5 M NaCl in PBST for 30 minutes and then rinsed two times with PBST. Samples then were incubated with Alexa-conjugated secondary antibodies (Invitrogen) diluted 1:1000 in block solution for one hour at room temperature. Slides were rinsed twice and washed three times for 5 minutes with PBST, incubated with Hoechst (Invitrogen) diluted 1:4000 in PBS, followed by two more 5 minute PBS washes. Slides were mounted using SlowFade Diamond (Thermo Fisher) and imaged using a Zeiss LSM 880 laser scanning confocal microscope.

### Cell counting and morphometrics

Osteoblasts and cranial neural crest cells of immunostained samples were counted using Fiji (National Institutes of Health). GFP-positive, DsRed-positive and EdU-positive cells were manually located and counted through image stacks. BSR3 lengths were measured on alizarin red and alcian blue stained flat mounts or alizarin red stained *Tg(sp7:eGFP)* larvae using the measurement tool in Fiji. Cell counting and BSR3 length statistical comparisons used Student’s two-tailed *t*-tests.

### Whole mount in situ hybridization

Digoxigenin-11-dUTP (Roche) labeled riboprobes were synthesized from PCR products amplified from larval cDNA. Reverse primers included a T7 transcription promotor sequence. *hoxa9b* primers were adapted from a previous study (van der Velden et al., 2012). Probes against *ezh1* and *ezh2* corresponded to 3’ UTRs to prevent cross detection of the paralogs. Primers for riboprobe templates: *hoxa9b* 5’-CCCGTGGTCCAGCAGCAGTC-3’, 5’-TAATACGACTCACTATAGTGCACTCACCACTCCCAACC-3’, *ezh1* 5’-ACGACCCACAATACTTTGCACTTC-3’, 5’-TAATACGACTCACTATAGGGTACATGTGACAGTATAACCTGAACC-3’, *ezh2* 5’-CCATCTACCTCTCTGAACAAATGCCT-3’, 5’-TAATACGACTCACTATAGGGCACCTTGTATTTGGAGGCAACAAC-3’. DNase treated probes were precipitated with LiCl and reconstituted in RNase-free water. Whole mount in situ hybridization was performed as described (Akerberg et al., 2017).

### Alkaline phosphatase assay

AB.9 cells (ATCC^®^ CRL-2298™) (Paw & Zon, 1999) were seeded in six well plates and cultured in Dulbecco’s Modified Eagle’s Medium (DMEM) supplemented with fetal bovine serum (10%), 1X penicillin-streptomycin (Gibco) and either DMSO (0.1%) or 5 μM EPZ-6438 (Medchem express) diluted in DMSO. After two days, cells were rinsed twice with PBS and switched to differentiation medium (Minimum Essential Medium alpha, 0.1% fatty acid free bovine serum albumin, 5 mM β-glycerophosphate, 50 ug/ml ascorbic acid, 100 nM dexamethasone). DMSO or EPZ-6438 was added when media was changed and once more 48 hours after media change. On day 6, cells were washed once with PBS, and then fixed for 1 minute with 4% formaldehyde in PBS. Fixed cells were washed once with PBS and once more with PBS containing 0.1% Tween-20. Cells were washed once for 5 minutes in developing buffer (100 mM Tris pH 9.5, 100 mM NaCl, 10 mM MgCl_2_) and then developed in the dark with fresh developing buffer supplemented with 225 ug/ml nitro blue tetrazolium (NBT) (Promega) and 175 μg/ml 5-bromo-4-chloro-3-indolyl-phosphate (BCIP) (Promega). Reactions were stopped with three PBS rinses. Cells were fixed with 4% PFA in PBS for 5 minutes at room temperature and then washed three times for 5 minutes. Cells were imaged with a Leica M165 FC stereomicroscope.

## ACKNOWLEDGEMENTS

We thank A. Akerberg for supporting initial *ezh2^sa1199^* studies; M. Solis-Wheeler and D. Nguyen for technical support; J. Nichols and C. Kimmel for experimental advice and consultations; the University of Oregon AqACS Facility for zebrafish care; the University of Oregon zebrafish community for support; A. Delaurier and C. Kimmel for *Tg(sp7:EGFP)b1212, Tg(runx2b:GAL4),* and *Tg(uas:E2CRIMSON)b1229* fish; and the Stankunas lab for input.

## FOOTNOTES

### Author contributions

G. A. Y., S. S. and K. S. designed experiments; G. A. Y. and S. S. performed experiments; G. A. Y. and K. S. prepared and wrote the manuscript.

### Data and material availability

Requests for materials should be addressed to K. S.

