## Supplemental Material for "Zebrafish Polycomb repressive complex-2 critical roles are largely Ezh2- over Ezh1-driven and concentrate during early embryogenesis"

**Figures S1-S8**

**Supplemental Movies 1-2**

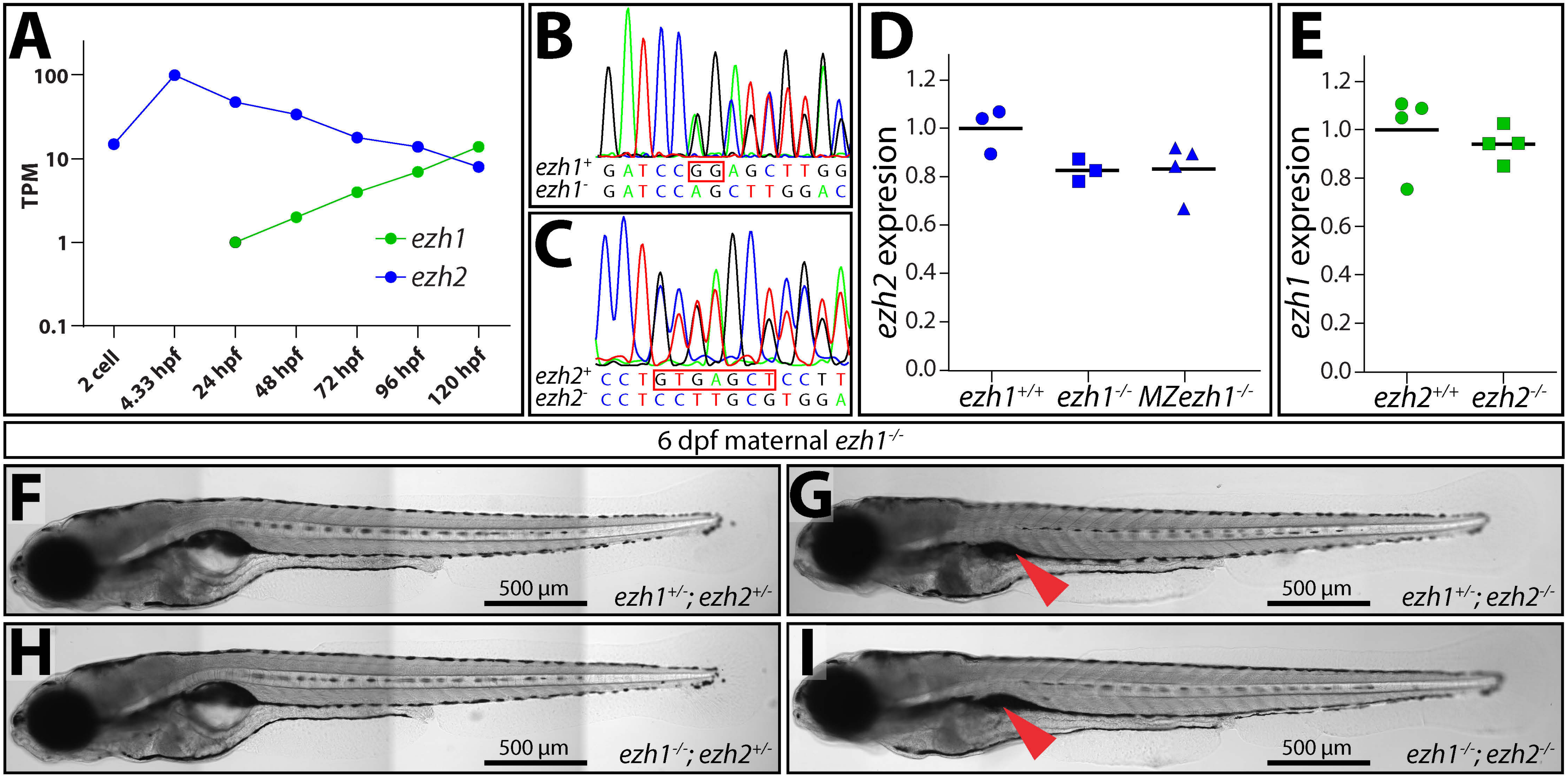


**Supplemental Figure 1**

**Figure S1. Maternal-zygotic *ezh1* mutants are viable and do not exacerbate zygotic *ezh2* loss-of-function overt defects. (A)** Graph showing *ezh1* (green) and *ezh2* (blue) transcript levels (transcript per million reads (TPM)) in 2-cell stage to 5 day post fertilization (dpf) wildtype zebrafish. Data is mined from a RNA-Seq developmental time course (White et al., 2017). **(B, C)** Chromatograms of sequenced amplicons from (B) *ezh1*$^{+/-}$ and (C) *ezh2*$^{+/-}$ larvae showing the CRISPR/Cas9-generated mutations in each respective allele (boxed in red). **(D, E)** Scatter plot graphs of normalized RT-qPCR results comparing (D) *ezh2* transcript levels in 3 dpf wildtype, *ezh1*$^{-/-}$ clutchmate, and *MZezh1*$^{-/-}$ larvae and (E) *ezh1* expression in 3 dpf wildtype versus *ezh2*$^{-/-}$ larvae. Neither allele produces compensatory expression of its corresponding paralog. **(F-I)** Whole mount differential interference contrast (DIC) microscopy images of 6 dpf maternal *ezh1* mutant siblings of indicated zygotic genotypes. Red arrowheads indicate under-inflated swim bladders of *ezh2*$^{-/-}$ larvae. Scale bars = 500 μm.

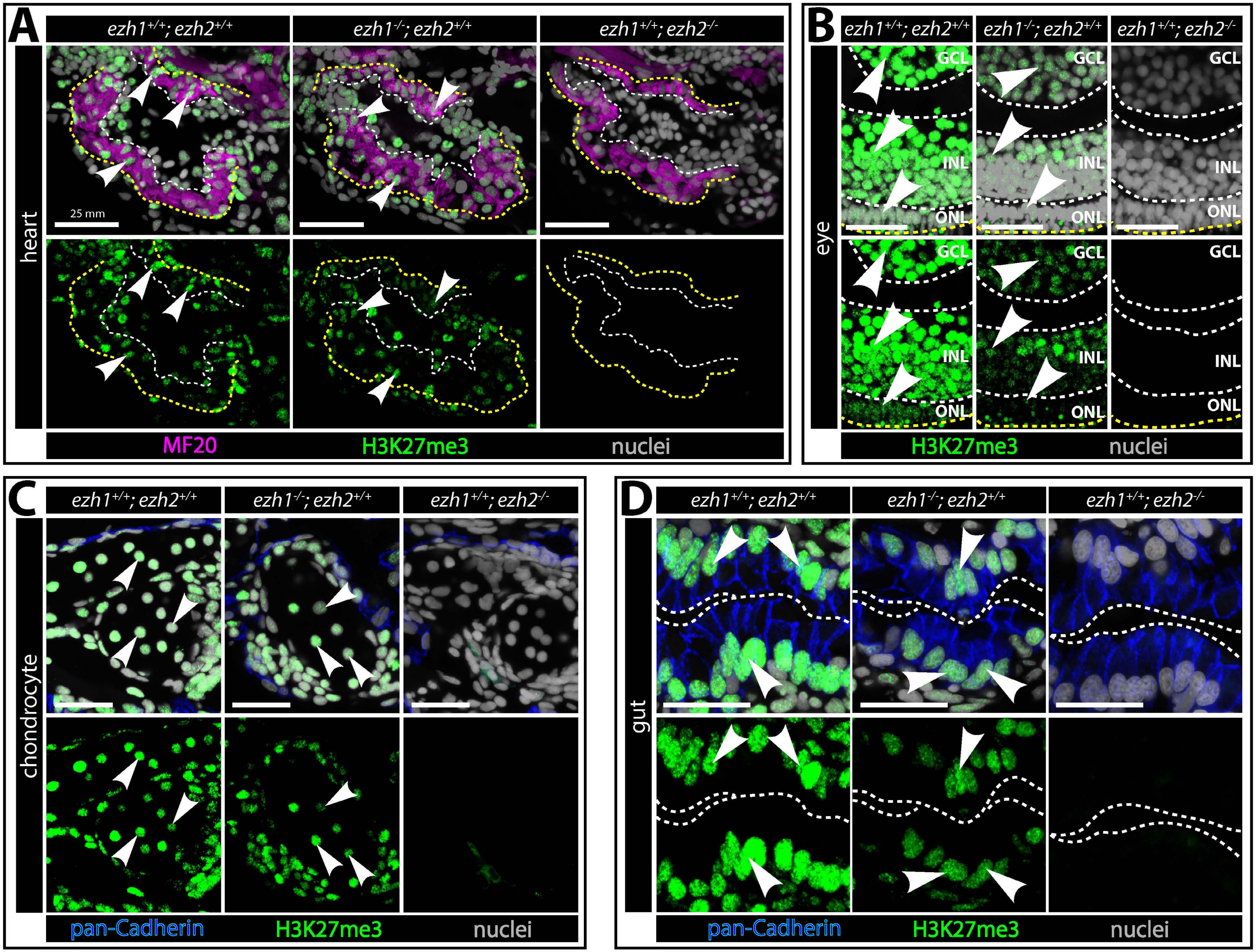


**Supplemental Figure 2**

**Figure S2. Ezh2 initiates while Ezh1 and Ezh2 dually maintain H3K27me3 in zebrafish larval organs**. **(A-D)** Maximum intensity projections of confocal imaged sectioned (A) heart, (B) eye, (C) craniofacial chondrocyte and (D) gut from 6 dpf wildtype, *ezh1*$^{-/-}$ and *ezh2*$^{-/-}$ larvae. As indicated, samples are immunostained with antibodies to detect myosin heavy chain (MF20, magenta), H3K27me3 (green) and pan-cadherin (blue). Nuclei are stained with Hoechst (grey). (A) Heart ventricles with surfaces and chambers outlined with dashed yellow and white lines, respectively. White arrowheads point to H3K27me3$^{+}$ cardiomyocytes. (B) White dashed lines separate ganglion cell layer (GCL), inner nucleated layer (INL) and outer nucleated layer (ONL) of the eye. Yellow dashed line outlines outer boundary of the eye. White arrowheads indicate H3K27me3$^{+}$ cells in each layer. (C) White arrowheads indicate H3K27me3$^{+}$ chondrocytes of the lower jaw. (D) Dashed white line marks the boundary between intestinal cells and intestinal lumen. White arrowheads point to H3K27me3$^{+}$ intestinal cells. Scale bars in all panels are 25 μm.

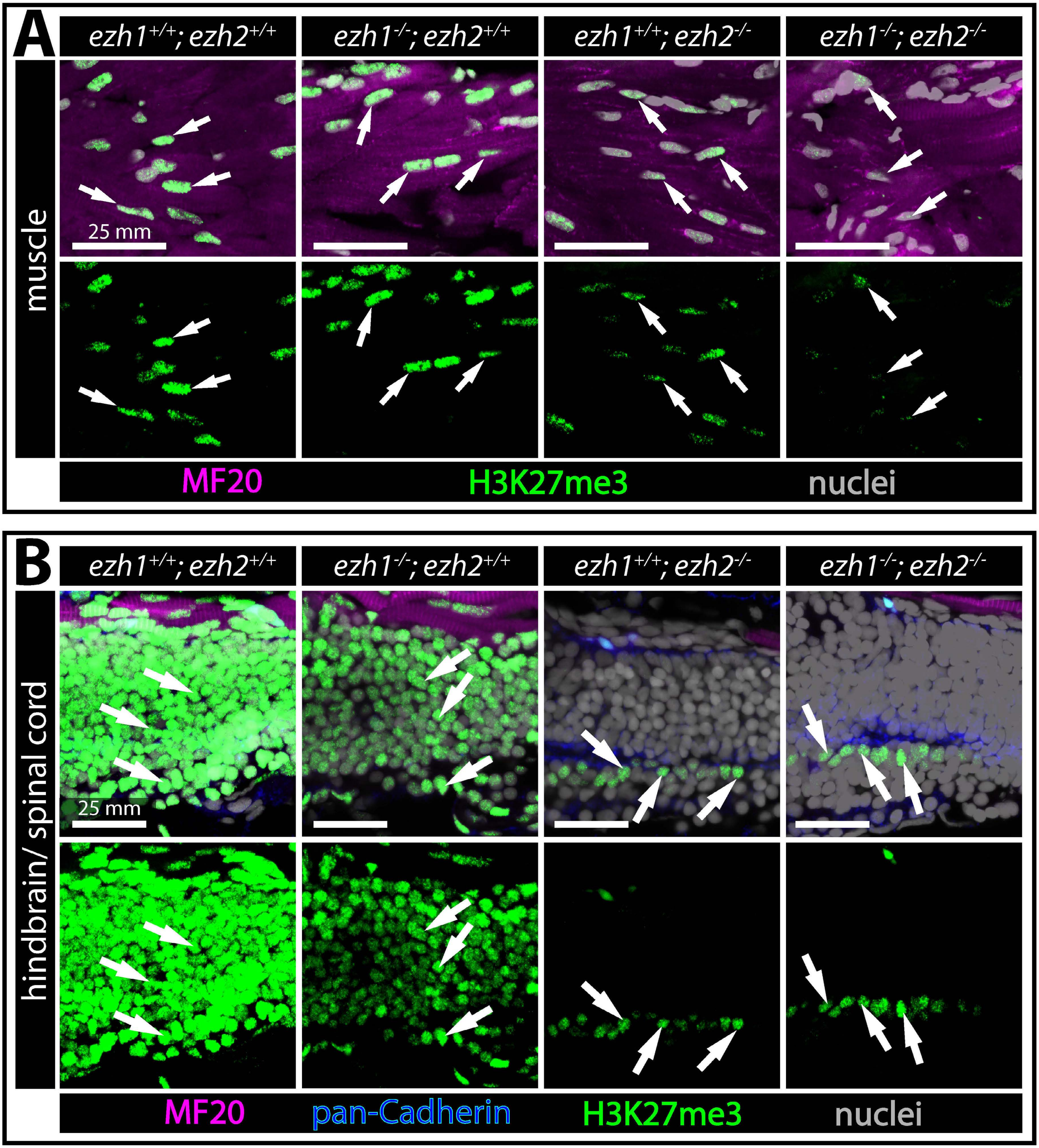


**Supplemental Figure 3**

**Figure S3. Residual H3K27me3 is retained in rare larval cells despite loss of zygotic H3K27me3 methyltransferases. (A, B)** Maximum projection confocal immunofluorescence images of wildtype, *ezh1*$^{-/-}$, *ezh2*$^{-/-}$ and *ezh1*$^{-/-}$; *ezh2*$^{-/-}$ sectioned (A) skeletal muscle and (B) nervous system tissue from 6 dpf larvae. (A) Sporadic skeletal muscle (MF20, magenta) cells retain low levels of H3K27me3 (green) even in the absence of zygotic Ezh1 and Ezh2 (far right panels). Similarly, H3K27me3 (green) is retained in a layer of nervous tissue cells (B) of *ezh1*$^{-/-}$; *ezh2*$^{-/-}$ larvae. Nuclei are Hoechst-stained (grey). White arrows highlight H3K27me3$^{+}$ cells. All scale bars are 25 µm.

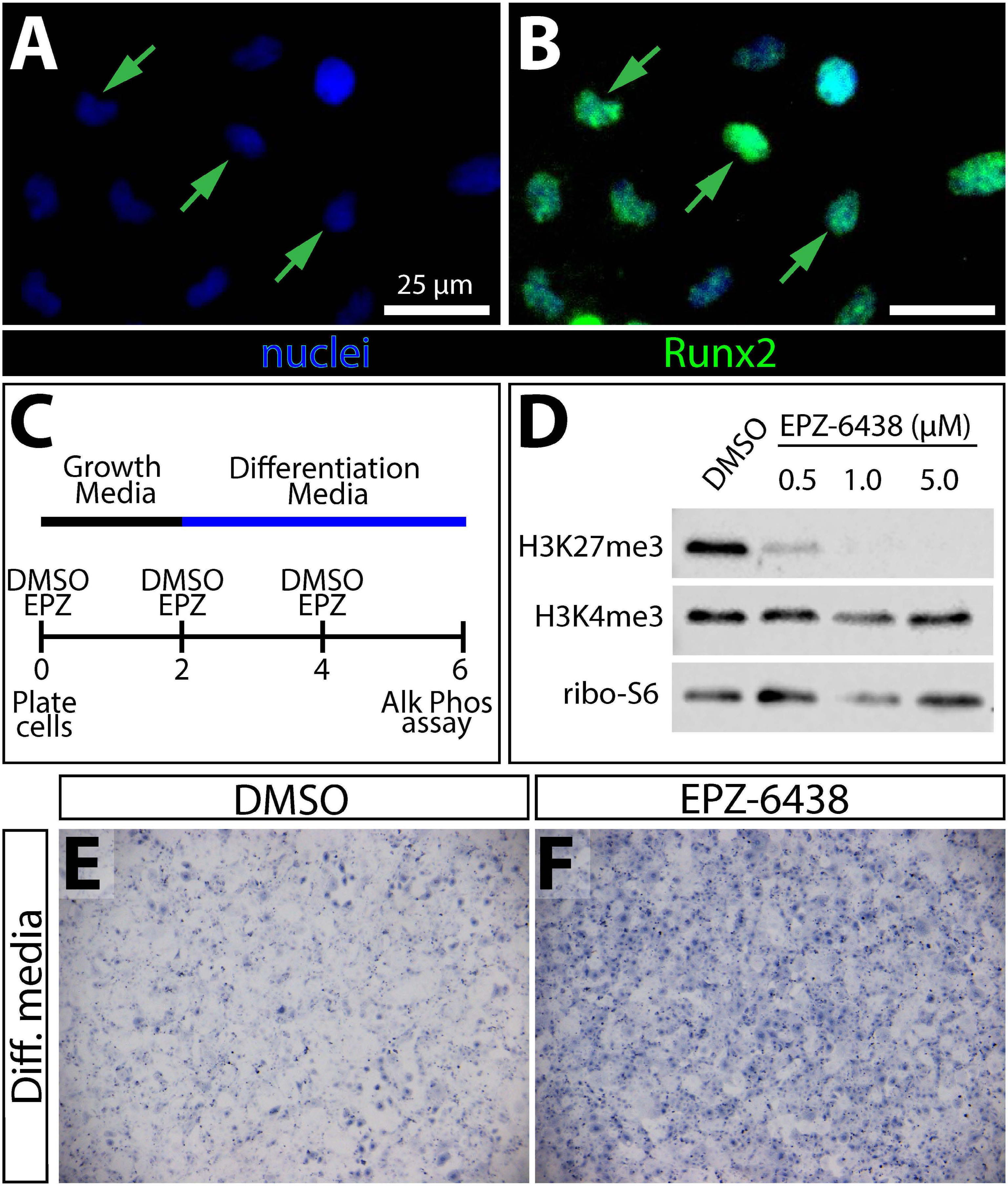


**Supplemental Figure 4**

**Figure S4. Ezh2 represses osteoblast maturation *in vitro*. (A, B)** Immunofluorescent staining of AB.9 cells using Runx2 (green) antibodies suggestive of their osteoblast identity. Green arrows point to Runx2$^{+}$ nuclei. Nuclei are stained with Hoechst (blue). Scale bars are 25 μm. **(C)** Diagram of treatment scheme to inhibit Ezh2 in AB.9 cells using the small molecule EPZ-6438 (Tazemetostat). AB.9 cells are plated and treated with either DMSO or EPZ-6438 for 48 hours in growth media. 48 hours after initial plating, cells are switched from growth to differentiation media and re-treated. EPZ-6438 is replenished at day 4 and then cells assayed for alkaline phosphatase activity, indicative of osteoblast differentiation, at 6 days post plating. **(D)** Immunoblots of whole cell lysates from AB.9 cells treated for four days with DMSO or varying concentrations of EPZ-6438 using antibodies against H3K27me3, H3K4me3, and ribosomal protein S6 (loading). **(E, F)** AB.9 cells treated with either DMSO or EPZ-6438 and assayed for alkaline phosphatase activity (blue).

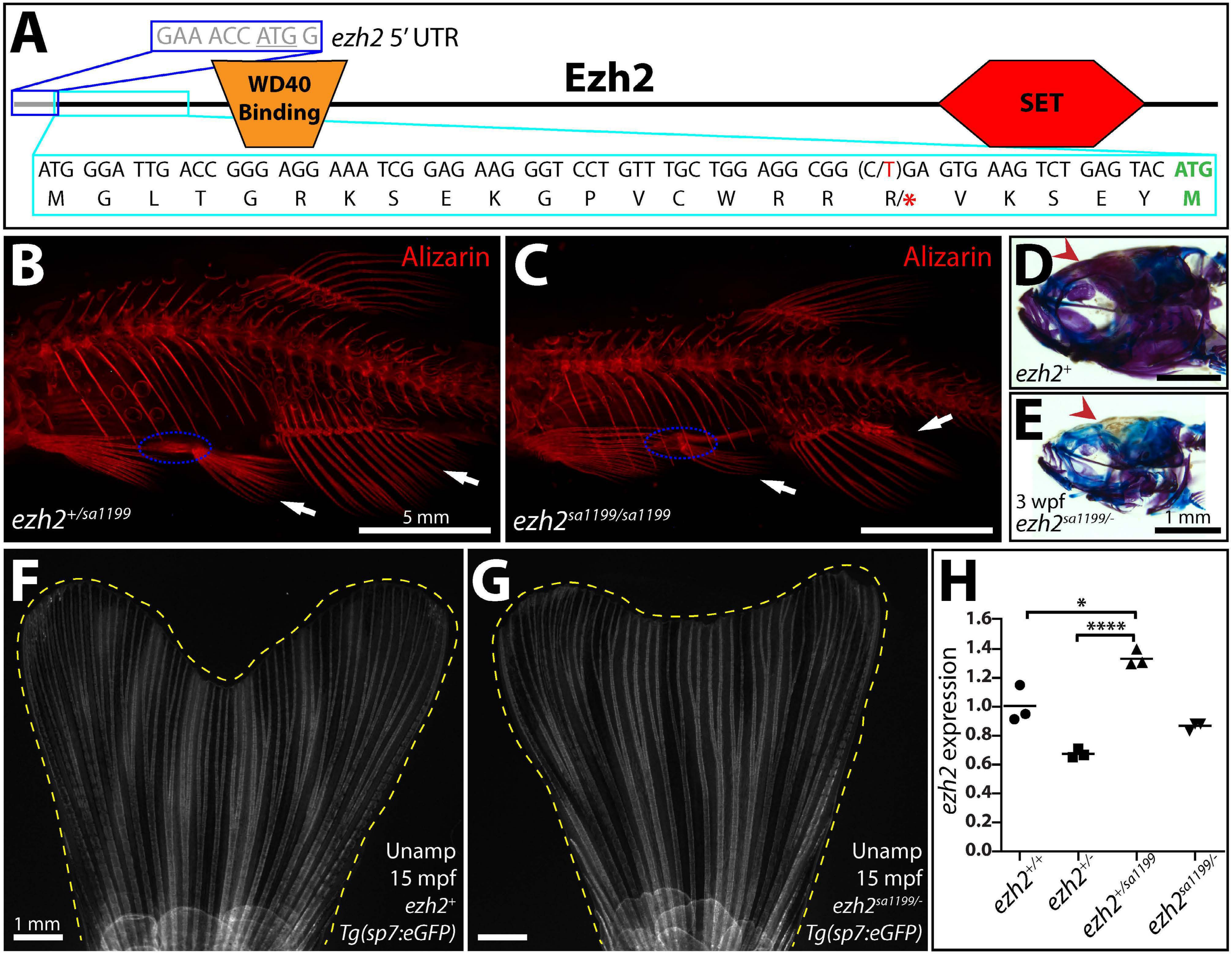


**Supplemental Figure 5**

**Figure S5. Hypomorph *ezh2*$^{sa1199}$ transcripts escape nonsense-mediated decay and produce truncated yet functional Ezh2 from an internal translation start site. (A)** Schematic of Ezh2 protein. The blue box and grey nucleotides show the *ezh2* start codon and immediately 5' untranslated region (UTR). The first 24 codons and amino acids are boxed in cyan. The *ezh2*$^{sa1199}$ point mutation is 52C>T (red), resulting in R18stop (red asterisk). A potential downstream in-frame translation start site at M24 (bold, green) would generate a truncated protein with all Ezh2 functional domains. **(B, C)** Skeletal preparations of five-month post fertilization control and *ezh2*$^{sa1199/sa1199}$ clutchmates stained with alizarin red (osteogenic bone, red). *ezh2*$^{sa1199/sa1199}$ adults have fewer bony rays in pelvic and anal fins (white arrows) and a reduced pelvic girdle (dashed blue oval). **(D, E)** Alcian blue (cartilage) and alizarin red (bone) stained heads of three-week old wildtype and *ezh2*$^{sa1199/-}$ clutchmates. Red arrowheads highlight reduced calcification of *ezh2*$^{sa1199/-}$ cranium, a phenotype consistent with fewer osteoblasts. **(F, G)** Whole mount fluorescent images of caudal fins from adult wildtype (F) and *ezh2*$^{sa1199/-}$ (G) *Tg(sp7:eGFP)* (bones, grey) clutchmates. Fins are outlined in yellow dashed line. **(H)** Scatter plot graph of RT-qPCR-determined *ezh2* transcript levels from adult fin regenerates of clutchmates of indicated genotypes. Expression is normalized to the mean of the *ezh2*$^{+/+}$ group after being first normalized to *rpl8* (reference gene). Fish with a copy of *ezh2*$^{sa1199}$ produce more *ezh2* transcripts, potentially enabling the survivability of *ezh2*$^{sa1199/-}$ and *ezh2*$^{sa1199/sa1199}$ fish in spite of minimal Ezh2 translation. Statistical significance determined by Student's two-tailed *t*-tests. * = P-value < 0.05, **** = P-value < 0.0001.

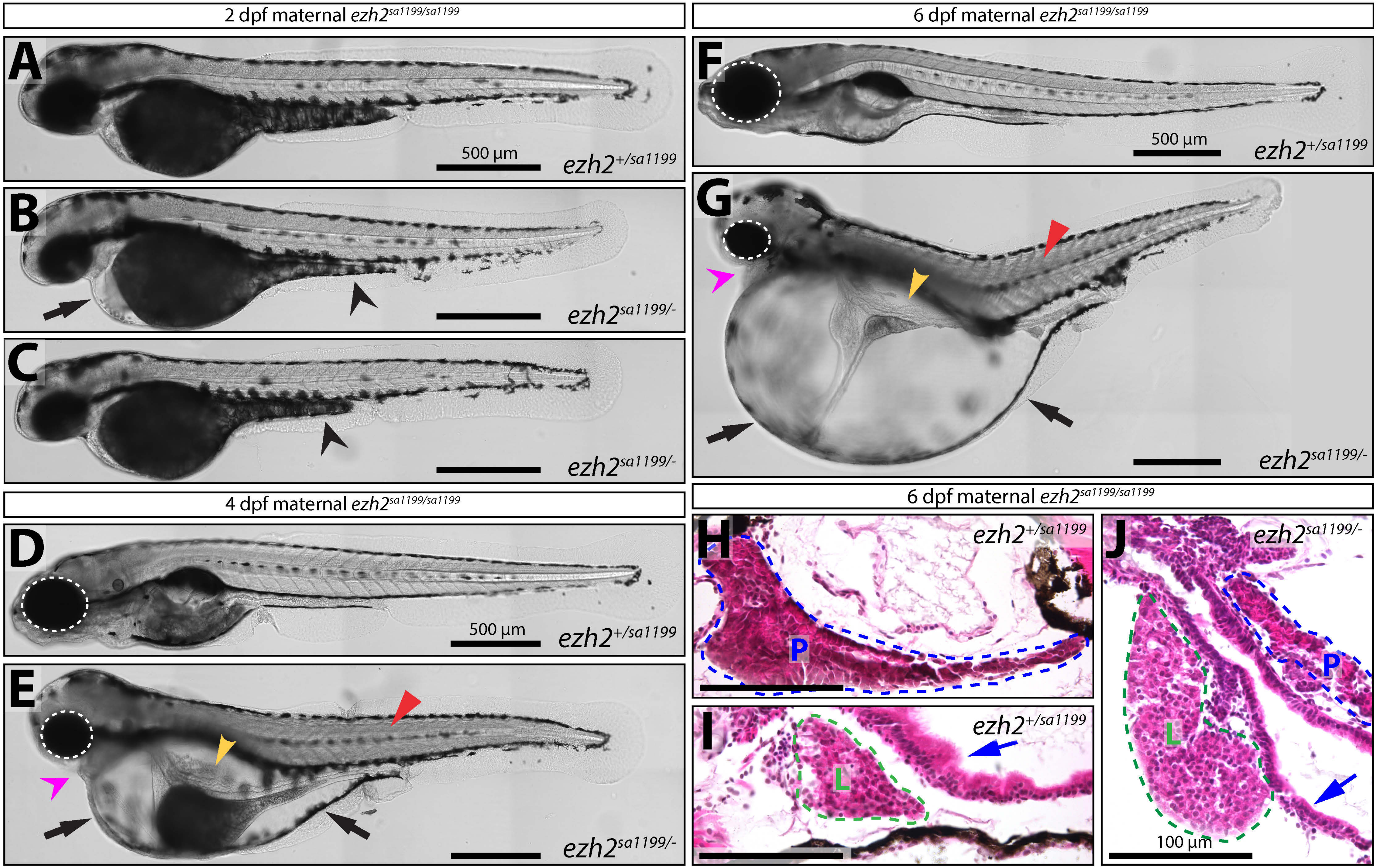


**Supplemental Figure 6**

**Figure S6. Combined maternal-zygotic *ezh2* has widespread organogenesis contributions. (A-G)** Whole mount DIC images of control and maternal-zygotic *ezh2* mutant clutchmate animals at (A-C) two dpf, (D, E) 4 dpf, and (F, G) 6 dpf. Black arrows point to edema first apparent around the heart at two dpf (B) and that progressively balloons with age (E, G). Black arrowheads indicate reduced yolk extension at two dpf (B, C). Yellow arrowheads mark the missing bulbous anterior intestine morphology in *Mezh2*$^{sa1199/sa1199}$*Zezh2*$^{sa1199/-}$ fish. Red arrowheads show abnormal muscle architecture. Magenta arrowheads indicate reduced jaws. Dashed white ovals emphasize differential eye sizes. Scale bars = 500 µm. **(H-J)** Hematoxylin and eosin stained paraffin sections of 6 dpf siblings of indicated genotypes from a ♀ *ezh2*$^{sa1199/sa1199}$ x ♂ *ezh2*$^{+/-}$ cross. The pancreas (H, J; outlined in blue dashed line, labeled with "P") and liver (I, J; outlined in green dashed line, labeled with "L") are abnormally shaped in maternal-zygotic *ezh2* mutants. The blue arrow indicates the poor columnar organization of intestinal epithelial cells in *Mezh2*$^{sa1199/sa1199}$*Zezh2*$^{sa1199/-}$ larvae. Scale bar is 100 µm.

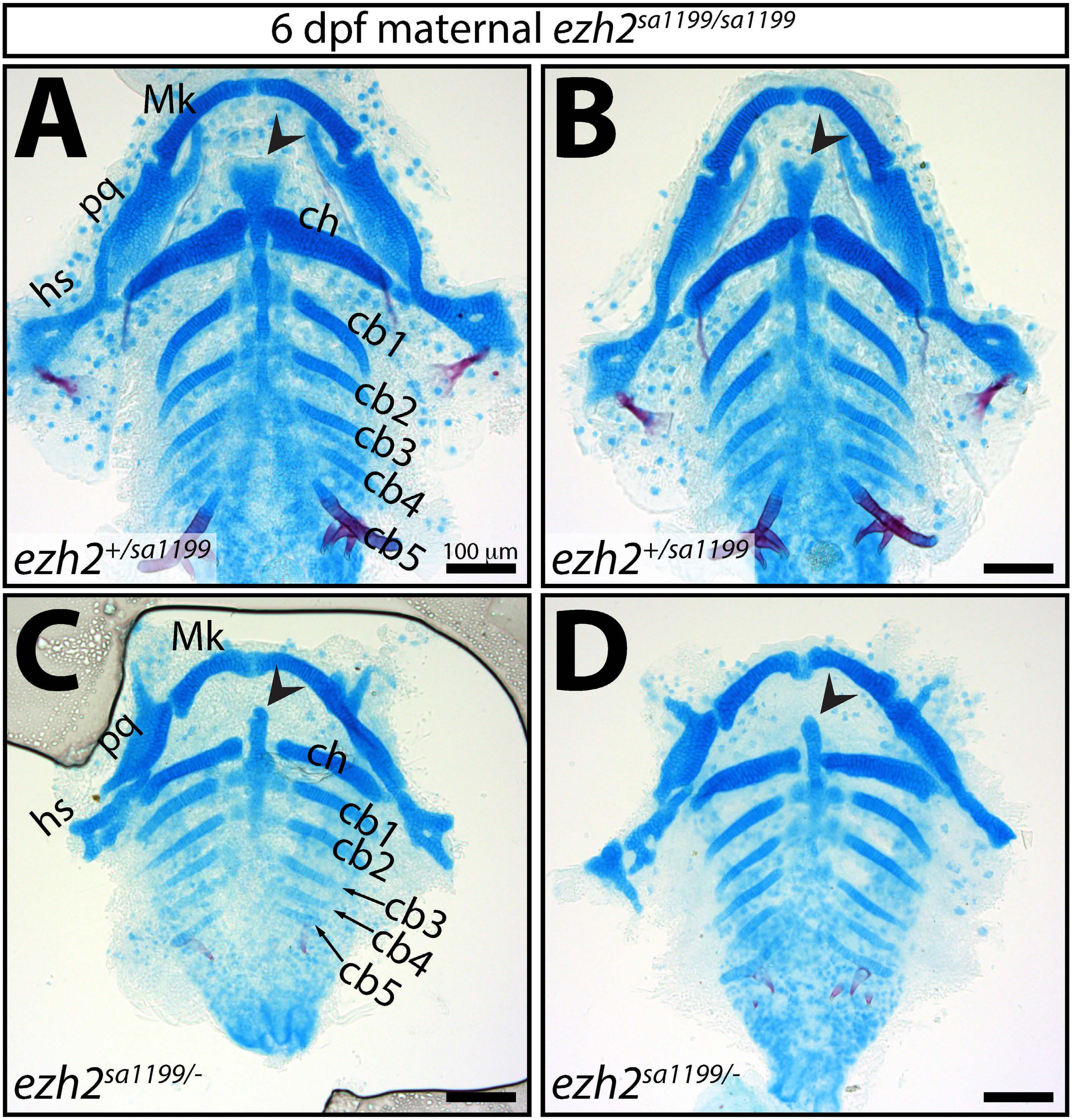


**Supplemental Figure 7**

**Figure S7. Maternal and zygotic *ezh2* promotes craniofacial skeletal size, shape and pattern**. **(A-D)** Ventral views of alcian blue (chondrocytes) and alizarin red (bone) stained viscerocranium from *Mezh2*$^{sa1199/sa1199}$ (A, B) *Zezh2*$^{+/sa1199}$ and (C, D) *Zezh2*$^{sa1199/-}$ 6 dpf clutchmate larvae. Ceratobranchial cartilage is diminished in maternal-zygotic *ezh2* mutants. Similarly, the basihyal (arrowheads) is smaller and fails to fan out in *Mezh2*$^{sa1199/sa1199}$*Zezh2*$^{sa1199/-}$ larvae. Abbreviations: Mk, Meckel's cartilage; pq, palatoquadrate; hs, hyosymplectic; ch, ceratohyal; cb, ceratobranchial. Scale bars are 100 µm.

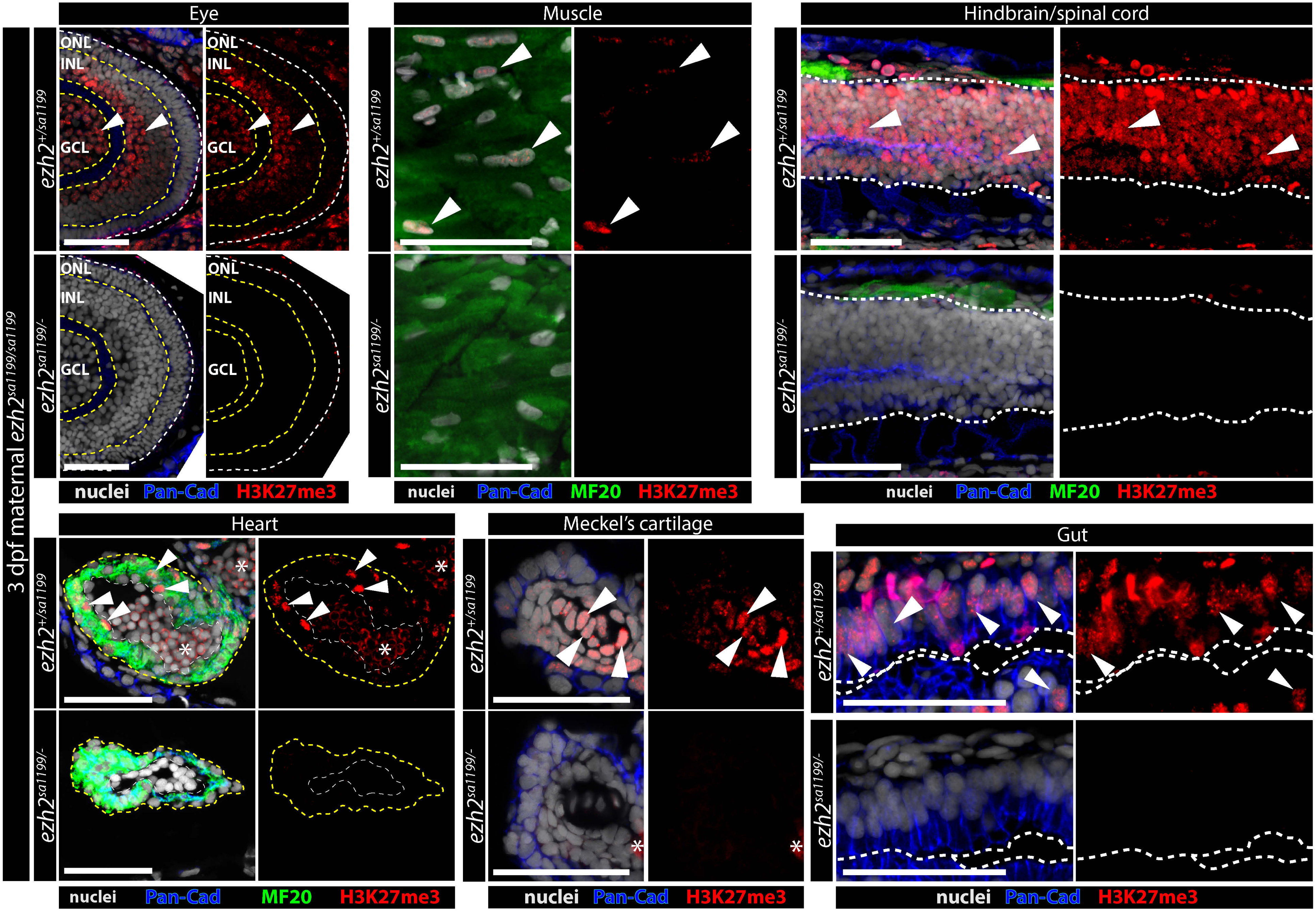


**Supplemental Figure 8**

**Figure S8.** $\boldsymbol{Mezh2^{sa1199/sa1199}Zezh2^{sa1199/-}}$ **embryos lack detectable bulk H3K27me3.** Confocal immunofluorescence images of sectioned eye, muscle, nervous tissue, heart, chondrocytes and gut from maternal zygotic *ezh2* mutant and zygotic control 3 dpf clutchmate embryos. Sections are stained for pan-Cadherin (blue), myosin heavy chain (green), and H3K27me3 (red), as indicated. Hoechst-stained nuclei are grey. A white dashed line outlines eye tissue. Yellow dashed lines separate the outer nucleated layer (ONL), inner nucleated layer (INL) and ganglion cell layer (GCL) of the eye. White dashed lines outline the hindbrain/spinal cord. Heart ventricles are outlined with yellow dashed lines. Ventricular chambers are traced with white dashed lines. White dashed lines separate intestinal lumen and epithelium. White arrowheads indicate H3K27me3$^+$ cells. Asterisks mark autofluorescence. All scale bars are 50 μm.

**Supplemental Movie 1. Zygotic Ezh2 promotes expansion of the opercle field without altering the onset of osteoblast differentiation.** Time lapse confocal imaging of wildtype (left) and *ezh2*$^{-/-}$ (right) clutchmate embryos carrying the *Tg(sp7:eGFP)* reporter (differentiating osteoblasts, grey) from 52 hpf until 72 hpf. GFP is detected at ~55 hpf in larvae of both genotypes. Scale bar is 50 µm.

**Supplemental Movie 2. Ezh2 promotes osteoblast migration for craniofacial morphogenesis.** Time lapse confocal imaging of wildtype (right) and *ezh2*-null (left) clutchmates monitored from 76 hpf until 96 hpf for osteoblast maturation and migration (*sp7:eGFP*, grey; weak, leaky expression in the ectoderm). Branchiostegal ray 3 (BSR3) osteoblasts appear stationary in *ezh2*$^{-/-}$ larvae compared to osteoblasts of wildtype larvae that migrate along the anteroposterior axis to expand the BSR3 field. Scale bar is 50 µm.
